# Inflammation primes the kidney for recovery by activating AZIN1 A-to-I editing

**DOI:** 10.1101/2023.11.09.566426

**Authors:** Segewkal Heruye, Jered Myslinski, Chao Zeng, Amy Zollman, Shinichi Makino, Azuma Nanamatsu, Quoseena Mir, Sarath Chandra Janga, Emma H Doud, Michael T Eadon, Bernhard Maier, Michiaki Hamada, Tuan M Tran, Pierre C Dagher, Takashi Hato

## Abstract

The progression of kidney disease varies among individuals, but a general methodology to quantify disease timelines is lacking. Particularly challenging is the task of determining the potential for recovery from acute kidney injury following various insults. Here, we report that quantitation of post-transcriptional adenosine-to-inosine (A-to-I) RNA editing offers a distinct genome-wide signature, enabling the delineation of disease trajectories in the kidney. A well-defined murine model of endotoxemia permitted the identification of the origin and extent of A-to-I editing, along with temporally discrete signatures of double-stranded RNA stress and Adenosine Deaminase isoform switching. We found that A-to-I editing of Antizyme Inhibitor 1 (AZIN1), a positive regulator of polyamine biosynthesis, serves as a particularly useful temporal landmark during endotoxemia. Our data indicate that AZIN1 A-to-I editing, triggered by preceding inflammation, primes the kidney and activates endogenous recovery mechanisms. By comparing genetically modified human cell lines and mice locked in either A-to-I edited or uneditable states, we uncovered that AZIN1 A-to-I editing not only enhances polyamine biosynthesis but also engages glycolysis and nicotinamide biosynthesis to drive the recovery phenotype. Our findings implicate that quantifying AZIN1 A-to-I editing could potentially identify individuals who have transitioned to an endogenous recovery phase. This phase would reflect their past inflammation and indicate their potential for future recovery.

## Introduction

The polyamines—namely putrescine, spermidine and spermine—are involved in a variety of fundamental biological processes such as transcription, translation, cell growth, differentiation, DNA repair, and aging (*1–3*). Polyamines are fully protonated at physiological pH and a substantial fraction of polyamines is associated with ribosomes (∼15%) and RNA (∼80%) (*4*). These nucleotide-bound polyamines facilitate global protein synthesis through their direct interaction with the translation machinery (*5, 6*). The critical role of polyamines in protein synthesis is further supported by the fact that cancer cells frequently exploit the polyamine pathway to enhance their growth (*7*). Conversely, polyamines are also essential for the activation of immune cells (*8, 9*), blurring the boundaries between therapeutic advantages and disadvantages.

The regulation of polyamine bioavailability is determined by a multitude of mechanisms including gut absorption, *de novo* synthesis, and the salvage pathways. In addition, polyamines significantly influence their own pathway through various post-transcriptional mechanisms (*1*). These mechanisms include ribosomal frameshifting (Ornithine Decarboxylase Antizyme 1), ribosomal occupancy of upstream open reading frames (Spermine Synthase and Spermidine/Spermine N1-acetyltransferase 1), stop codon readthrough (Adenosylmethionine Decarboxylase 1), posttranslational modification of Eukaryotic Translation Initiation Factor 5A (hypusination), as well as post-transcriptional mRNA editing of Antizyme Inhibitor 1 (AZIN1) from adenosine to inosine (A-to-I). This A-to-I editing results in a non-synonymous amino acid mutation, as inosines are translated as guanosines (*10*). The presence of these intricate regulatory mechanisms within this pathway underscores the crucial importance of controlling polyamine levels in response to various environmental stresses.

The kidney is an organ with exceptionally high metabolic demands (*11*), making it susceptible to various stressors such as diabetes and sepsis, which can disrupt polyamine homeostasis. Indeed, a recent study has highlighted that altered polyamine metabolism is a unifying feature across more than 10 different kidney injury models in mice, as well as in the context of post-kidney transplantation in humans (*12, 13*). Although the importance of polyamines in kidney biology is indisputable, their exact role under stress conditions remains unclear. The supplementation of polyamines and the modulation of the polyamine pathway have yielded diverse outcomes in multiple models of kidney injury, ranging from providing protection to exacerbating tissue damage (*14–21*). These varying results underscore the need for a more systematic examination of the roles of polyamines across specific disease timelines and trajectories.

Defining timelines and stages of any kidney disease is highly challenging. Due to variations in disease progression among patients, a uniform physical timescale cannot be universally applied. We reasoned that the precisely controlled, stepwise reactions embedded in the polyamine pathway could serve as the basis for constructing a molecular clock. This path of investigation has led to our present findings, which demonstrate that AZIN1 A-to-I editing is strikingly prevalent and occurs at specific points along disease timelines in both mouse models and humans. As such, AZIN1 A-to-I editing can serve as a molecular clock to stage various forms of kidney disease.

AZIN1 is a key regulatory enzyme controlling the upstream entry point to the polyamine pathway (*22*). The A-to-I editing of AZIN1 confers a gain-of-function phenotype in several forms of cancer, contributing to aggressive tumor behavior (*23–26*). The role of AZIN1 editing is also implicated in hematopoietic stem cell differentiation (*27*). Most recently, transient AZIN1 editing has been reported in cases of COVID-19 infection (*28*). However, the clinical implications of AZIN1 editing in non-cancerous kidney diseases remain unclear.

By combining a series of sequencing and genetic approaches, we found that AZIN1 edited state confers an advantage over the unedited state by upregulating the polyamine pathway and co-opting glycolysis and nicotinamide biosynthesis, culminating in a metabolically robust phenotype. Using an extensively characterized murine model of endotoxemia, we also provide a genome-wide, time-anchored map of A-to-I editing, serving as a novel framework for the development of molecular staging in kidney disease.

## Results

### AZIN1 A-to-I editing is widespread in non-cancerous conditions

Using a model of endotoxin preconditioning, we have previously identified that increased polyamine levels are a key feature of the robust protective phenotype against severe sepsis (*14*). Increases in polyamine levels are also reported by others during the recovery phase of ischemia-reperfusion injury (*29*). Conversely, inhibiting a branch of polyamine pathway can also lead to tissue protection against multiple models of kidney diseases (e.g., inhibition of eIF5A hypusination or Ornithine decarboxylase) (*15, 16, 30–32*). These contrasting findings suggest that the role of polyamines is context dependent, such as the severity of tissue injury or timing of intervention. To understand the role of polyamines broadly in various stress conditions, here we first interrogated a large clinical dataset in which stranded RNA-seq was performed on whole blood collected from children before and after contracting malaria (*33*). Through prospective surveillance, the patients were categorized into 1) early fever (infection with concurrent fever), 2) delayed fever (infection with a delay of 2-14 days until development of fever), and 3) immune (infection without progression to fever). We found that AZIN1 A-to-I editing at chromosome 8:102829408 (hg38), a known A-to-I editing site (*34*), was highly prevalent in this cohort, albeit at different time points among the three groups (**Fig 1A, B, Suppl Fig 1A-C**). Notably, children in the early fever group had low levels of AZIN A-to-I editing at baseline but showed an increase in editing after malaria infection. In contrast, children in the delayed and immune groups exhibited surprisingly high levels of A-to-I editing at baseline that were sustained over time. This raises the possibility that AZIN1 A-to-I editing early in the course of malaria infection could have a beneficial role in controlling disease progression.

**Figure 1.**
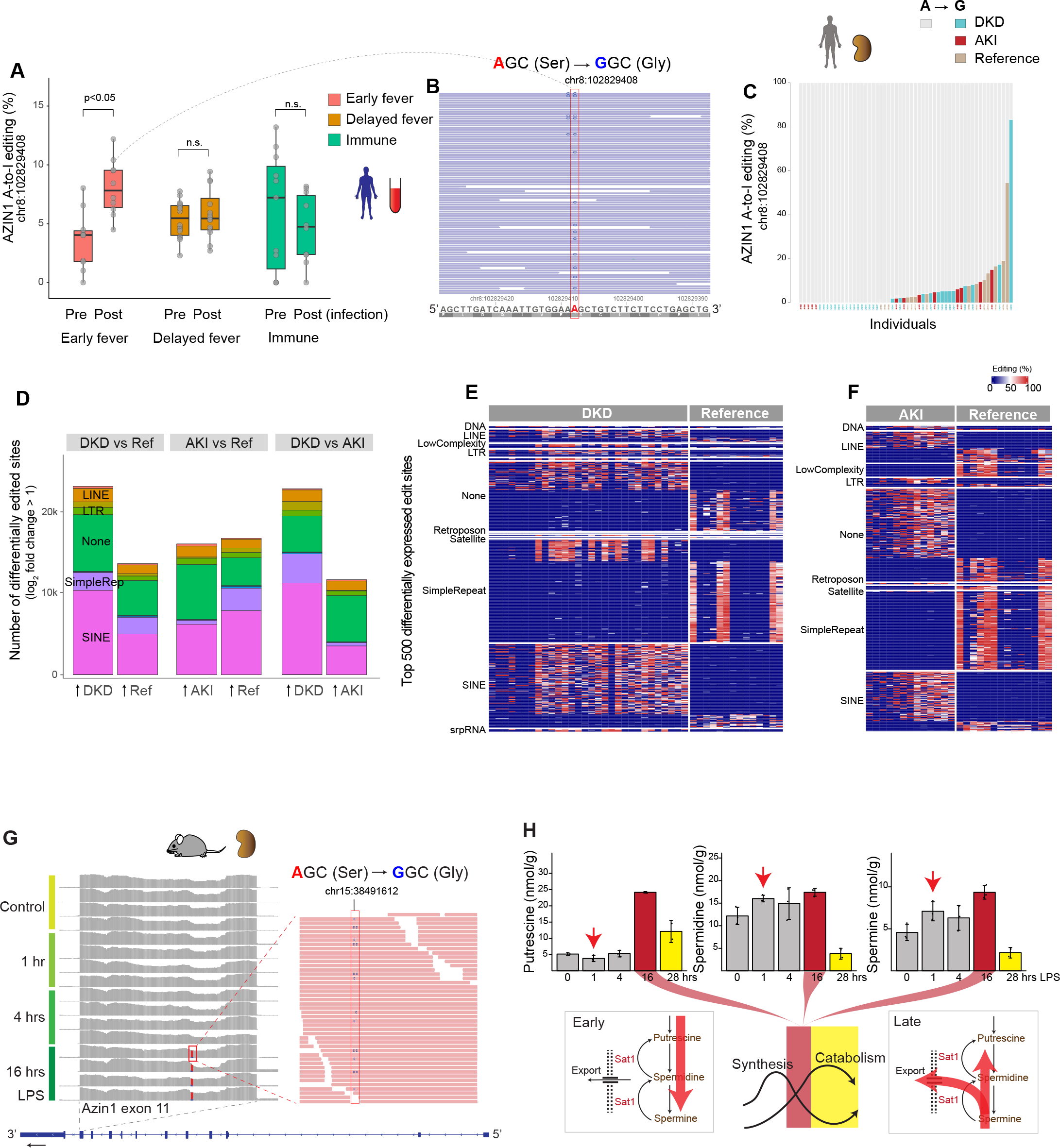
AZIN1 A-to-I editing status in non-cancerous diseases in humans and mice. (**A**) Distribution of AZIN1 A-to-I editing rates (% of edited reads over total reads) in prospectively collected blood from male children aged 6 to 11 years, before and after *Plasmodium falciparum* malaria infection. Individuals were classified as early fever (symptomatic and first-time infection), delayed fever (asymptomatic and first-time infection, subsequently developing malarial symptoms), and immune (infected but never developed symptoms). (**B**) Representative read coverage near the AZIN1 editing site for one sample. Note that inosine is sequenced as guanosine. The human AZIN1 gene is encoded on the minus strand, hence the T-to-C mutation, not A-to-G, in the coverage track. Light blue-colored reads (F2R1 paired-end orientation) indicate the proper directionality of reads mapped to the minus strand. (**C**) Distribution of AZIN1 A-to-I editing rates in kidney biopsies with a pathology diagnosis of diabetic kidney disease (DKD), acute kidney injury (AKI), or reference nephrectomy samples. Each column represents one sample. (**D**) Stacked bar chart summarizing total numbers of differentially expressed A-to-I editing sites genome-wide under the indicated conditions. For each comparison, editing sites are divided on the x-axis based on the direction of fold change. For example, in the DKD vs reference comparison, approximately 20K sites are more edited in DKD, whereas approximately 10K sites are more edited in reference nephrectomy samples. (**E**) Heatmap displaying the top 500 differentially expressed A-to-I editing sites between diabetic nephropathy and reference nephrectomy samples. The differentially expressed sites are categorized based on repeat classes. (**F**) Comparison between AKI biopsies and reference nephrectomy samples. (**G**) Total RNA-seq read coverage around the Azin1 A-to-I editing site in the mouse kidney tissues under the indicated conditions. A magnified view at 16 hours after LPS is shown for one sample. The mean editing rates are 1.8%, 1.8%, 1.2%, and 19.0 % at 0, 1, 4, and 16 hours, respectively. (**H**) Kidney tissue polyamine levels as determined by targeted metabolomics under the indicated conditions.

Next, we interrogated stranded RNA-seq data of human kidney biopsies obtained from our biobank and the Kidney Precision Medicine Project (*35, 36*). We found that AZIN1 editing is common in non-cancerous kidney tissues, including those with diabetic kidney disease, acute kidney injury (AKI), and even reference nephrectomy (**Fig 1C**). However, no difference was found in the extent of AZIN1 editing among the three groups. This may be due to the fact that these biopsies were obtained at various stages in the diabetes and AKI timelines (**Suppl Fig 1D**). Similarly, the reference biopsies are known to sustain variable degrees of ischemic injury, thus exhibiting some AKI phenotype. In addition, reference nephrectomy samples were derived from tissues adjacent to renal cell carcinoma, which may also influence AZIN1 A-to-I editing status. Nevertheless, genome-wide examination did reveal significant differences among the three groups in A-to-I editing at tens of thousands of sites (**Fig 1D**; see Methods). Overall, diabetic kidneys showed more extensive genome-wide A-to-I editing than nephrectomies and AKI samples. When focusing on the top differentially edited sites, reference nephrectomy samples had A-to-I editing predominantly within simple repeat regions, whereas AKI and diabetic samples had A-to-I editing within short interspersed nuclear elements (SINE, such as Alu elements; **Fig 1E-F**). The differential editing in transposable elements such as SINE may have profound implications on disease unfolding (*37*).

### Changes in AZIN1 A-to-I editing and polyamine metabolism across the sepsis timeline

To understand the role of AZIN1 editing and polyamine metabolism in the kidney, we next interrogated a well-characterized animal model of endotoxemia. In this specific model, the kidney goes through precise stages, starting with classic NFκB-mediated acute inflammation, followed by interferon responses and the integrated stress response, and culminating in metabolic and translation shutdown (*38–40*). Single-cell RNA sequencing revealed that Azin1 is expressed in all cell types in the kidney (**Suppl Fig 2A**). Furthermore, Ribo-seq analysis showed that Azin1 gene expression and its translation efficiency remained constant throughout the course of endotoxemia (**Suppl Fig 2B**). However, we found that Azin1 A-to-I editing status varied significantly over the same time period (**Fig 1G**). While the extent of A-to-I editing was minimal at baseline and during the early phases of endotoxemia, it significantly increased during the later stages of sepsis in this model. In fact, we observed a consistent and robust increase in Azin1 A-to-I editing at around 16 hours and later time points after endotoxin exposure. We have previously shown that this 16-hour time point corresponds to a critical transition phase between translation shutdown and subsequent tissue recovery (*38, 39*). Thus, editing of Azin1 at this precise time point may serve as a clock to stage endotoxemia. Furthermore, since Azin1 A-to-I editing confers a gain-of-function, it may also signal a change in polyamine metabolism that aids tissue healing.

While polyamine metabolism could be cell-type specific (**Suppl Fig 2C, D**), quantitation of polyamine levels in bulk kidney tissues revealed marked changes throughout the sepsis timeline (**Fig 1H**). One hour after endotoxin challenge, tissue spermidine and spermine levels increased, while putrescine levels decreased. This is indicative of increased flux along the polyamine conversion axis (red arrows; **Fig 1H**). This increase of polyamines may be a participant in the early inflammatory response. At time points beyond 16 hours, we noted a significant depletion of polyamines. This depletion paralleled underlying changes in key enzymes involved in polyamine metabolism (**Suppl Fig 2E, F**). For example, the expression of Ornithine Decarboxylase 1 and other polyamine anabolic genes decreased, which may explain the depletion of polyamines and translation shutdown in this late phase of sepsis. Because Azin1 is an activator of Ornithine Decarboxylase 1, its gain-of-function conferred by A-to-I editing may be essential for preventing irreversible tissue shutdown.

### AZIN1 A-to-I uneditable cells are compromised upon nutrient deprivation and mitochondrial inhibition

To elucidate the functional significance of AZIN1 editing, we next designed two homozygous clonal cell lines using the CRISPR knock-in strategy (**Fig 2A, Suppl Fig 3A**). The first line contains a constitutively edited AZIN1, resulting in an A-to-I locked state (AGC serine to GGC glycine). The second line is an A-to-I uneditable variant in which the editing site is disrupted while preserving the codon composition (AGC serine to TCC serine). A-to-I locked or uneditable state did not lead to changes in the abundance or stability of the AZIN1 protein (**Fig 2B, C**). We found that A-to-I locked cells exhibit accelerated cell growth compared to wild-type and A-to-I uneditable cells, all of which share an otherwise identical genetic background (HEK293T; **Fig 2D, E, Suppl Fig 3B, C**). The level of A-to-I editing in the wild-type cells was minimal (∼0%). However, the growth curve of the wild-type cells fell between those of the A-to-I locked and uneditable cells. This suggests that transient and low grade AZIN1 editing is operative under normal condition, contributing to healthy cellular growth. In support of the rapid growth rate observed in the A-to-I edited state, multiple genes involved in cell growth and differentiation were upregulated in the A-to-I locked cell line (e.g., BMP2/Bone Morphogenetic Protein 2, IGFBPL1/Insulin Like Growth Factor Binding Protein Like 1, PGF/Placental Growth Factor; **Fig 2F, Suppl Fig 3E**, https://connect.posit.iu.edu/azin1/).

**Figure 2:**
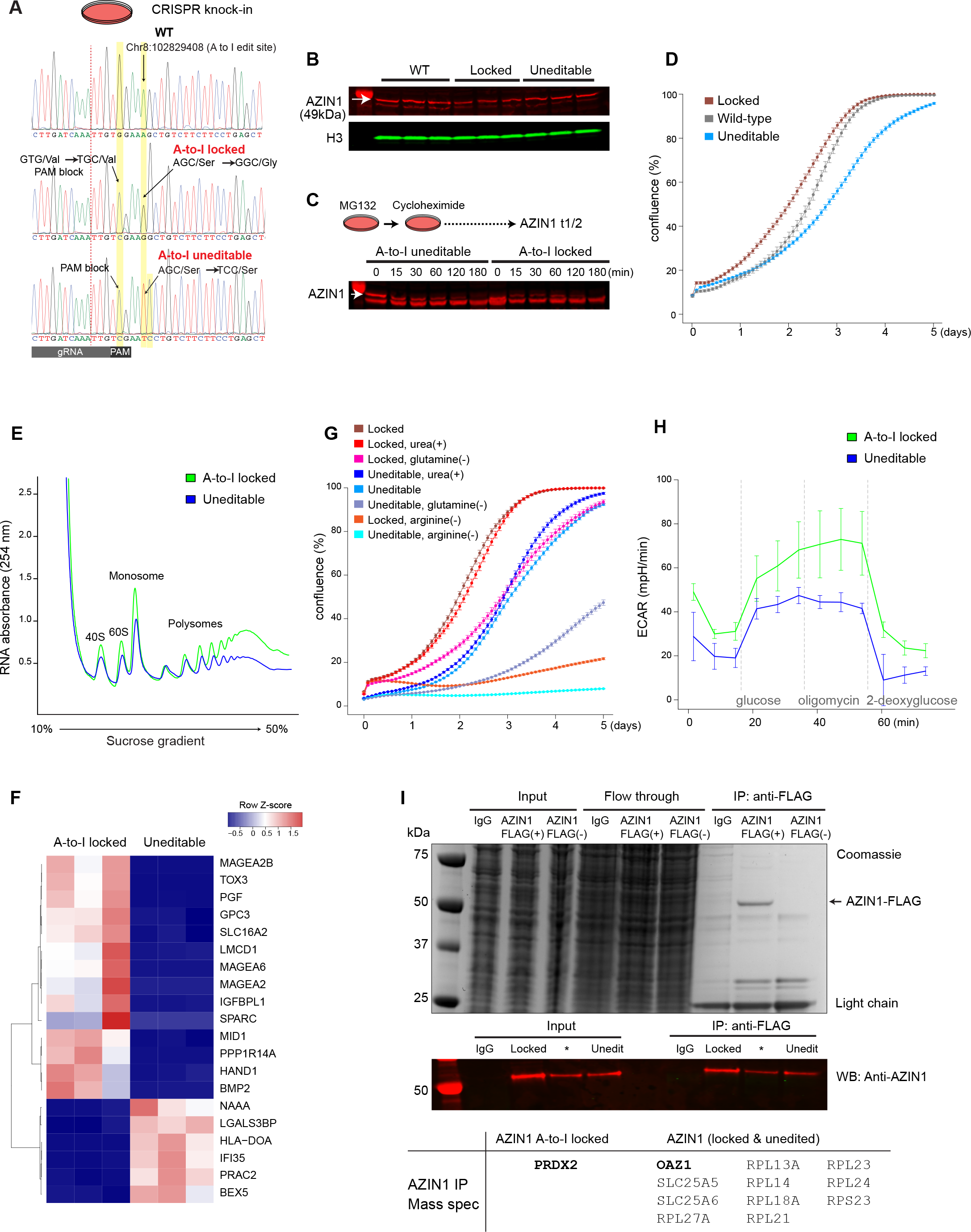
Azin1 A-to-I uneditable state hinders cell growth and limits glycolytic capacity. (**A**) Sanger sequencing chromatograms for wild-type (HEK293T; top), AZIN1 A-to-I locked (middle), and AZIN1 A-to-I uneditable homozygous cell lines (bottom). Homology-directed repair donor oligos used for CRISPR knock-in are shown in **Supplemental Figure 3A**. (**B**) Western blotting for AZIN1 under the indicated conditions (approximately 70% confluency). (**C**) Determination of AZIN1 protein turnover under the indicated conditions. Nascent protein synthesis was inhibited with cycloheximide 250 µg/ml. The arrow points to AZIN1. The bands below AZIN1 result from the inhibition of proteasomal degradation with MG132. n=2 biological replicates. (**D**) Real-time monitoring of cell growth for AZIN1 A-to-I locked, uneditable, and wild-type cells (n=3 independent experiments with n=6 technical replicates for each experiment). Representative images are shown in **Supplemental Figure 3C**. (**E**) Polyribosome profiling of AZIN1 A-to-I locked and uneditable cell lines. n=3 independent experiments. Mean polysome-to-monosome ratios for A-to-I locked and uneditable genotypes are 4.1 and 3.6, respectively. (**F**) Heatmap of the top 20 differentially genes between AZIN1 A-to-I locked and uneditable cell lines as determined by RNA-seq (https://connect.posit.iu.edu/azin1/). (**G**) Cell growth under the indicated conditions. Representative images are shown in **Supplemental Figure 3D**. (**H**) Extracellular acidification rates under the indicated conditions (Seahorse glycolysis stress test). n=3 independent experiments with n=3 technical replicates for each experiment. (**I**) Identification of AZIN1-interacting molecules by mass spectrometry. The top panel shows Coomassie staining for input, flow through, and immunoprecipitated unfractionated lysates from IgG control and transfection of FLAG-tagged AZIN1 or AZIN1 without FLAG plasmids. The lower panel shows Western blotting for AZIN1. Cells overexpressing FLAG-tagged A-to-I locked AZIN1 or uneditable plasmids were fractionated into cytoplasmic and nuclear compartments and immunoprecipitated using the anti-FLAG antibody (cytoplasmic fraction is shown. See also **Supplemental Figure 3G**). Summary of co-precipitated proteins with AZIN1 is presented in the bottom table. n=3 independent experiments. *denotes a plasmid construct not used in this manuscript.

As expected, the depletion of arginine exhibited a profound growth inhibitory effect on cell proliferation, which was more notable in the uneditable cell line (**Fig 2G; Suppl Fig 3D**). Conversely, the supplementation of urea, known to enhance polyamine biosynthesis (*41, 42*), rescued cell proliferation in the uneditable cell line. This effect was not observed in the A-to-I locked cell line, suggesting that polyamine synthesis is maximized in the A-to-I locked state. In addition, the impact of glutamine depletion was less pronounced in the A-to-I locked cell line (**Fig 2G**). Surprisingly, glycolysis stress test revealed marked differences in extracellular acidification rates between A-to-I locked and uneditable cell lines. Specifically, the uneditable cell line lacked a compensatory glycolytic response upon ATP synthase inhibition (**Fig 2H, Suppl Fig 3F, G**). While the exact mechanism remains uncertain, these findings offer a new perspective on the involvement of AZIN1 A-to-I editing in metabolic flexibility. This is especially pertinent in situations such as cancer and ischemia-reperfusion injury. In this regard, non-polyamine related functions of A-to-I locked AZIN1 cannot be excluded. For example, immunoprecipitation of AZIN1 identified that A-to-I edited AZIN1 uniquely binds to the thiol-specific peroxidase Peroxiredoxin2 (**Fig 2I, Suppl Fig 3H**).

### Azin1 A-to-I editing confers resilience through the orchestration of multiple protective pathways

To gain further insight into the *in vivo* implications of Azin1 A-to-I editing, we created two CRISPR knock-in mouse models, representing both A-to-I locked and uneditable states (**Fig 3A, Suppl Fig 4A**). Because Azin1 edited status had a significant effect on glycolysis, we examined its role in an ischemia-reperfusion model of kidney injury. We found that Azin1 locked mice had less severe kidney damage after ischemia-reperfusion injury as compared to the uneditable mice (**Fig 3B**). In addition to the reduction in serum creatinine levels, the less pronounced tissue damage in A-to-I locked mice was reflected by the better-preserved global translation (**Fig 3C**). Note that no discernible difference was observed in eIF5A hypusination between the two knock-in models (**Fig 3D**). This suggests that the beneficial effects of Azin1 A-to-I editing on translation are mediated through hypusination-independent polyamine pathways.

**Figure 3:**
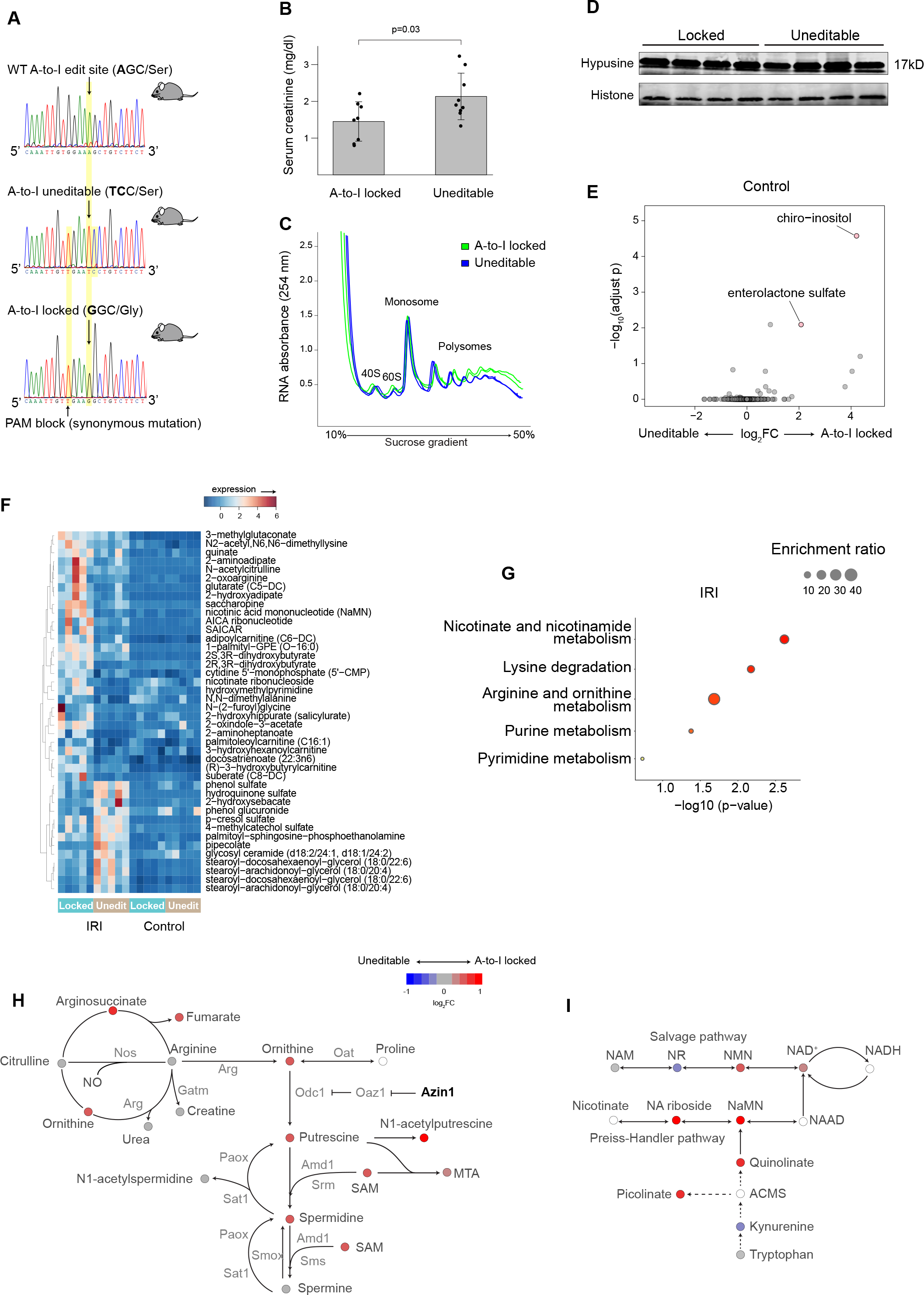
Azin1 A-to-I locked state limits kidney injury by upregulating polyamines and other protective pathways. (**A**) Sanger sequencing chromatograms for wild-type (top), Azin1 A-to-I uneditable (middle), and Azin1 A-to-I locked homozygous mice (bottom). The CRISPR knock-in strategy is depicted in **Supplemental Figure 4A**. (**B**) Serum creatinine levels 24 hours after a 20-minute bilateral ischemia-reperfusion injury. (**C**) Polyribosome profiling of kidneys from Azin1 A-to-I locked and uneditable mice 24 hours after ischemia-reperfusion injury. Two representative biological replicates are shown for each genotype. Mean polysome-to-monosome ratios for A-to-I locked and uneditable genotypes are 3.3 and 2.8, respectively. (**D**) Western blotting for hypusine in the kidney after ischemia-reperfusion injury. (**E**) Volcano plot showing the top two differentially expressed metabolites. The x-axis depicts the log2 fold change of A-to-I locked-to-uneditable ratio, and the y-axis depicts −log10 adjusted p values. Global untargeted metabolomics, n=5 for each condition. (**F**) Heatmap displaying the top differentially expressed metabolites between Azin1 locked and uneditable mice after ischemia-reperfusion injury (adjusted p < 0.05 for all listed metabolites). (**G**) Pathway enrichment analysis of differentially expressed metabolites between Azin1 A-to-I locked and uneditable mouse kidneys after ischemia-reperfusion injury. (**H**) Metabolite ratios (log2 fold change of A-to-I locked-to-uneditable) mapped to the polyamine pathway and pseudo-colored according to the indicated scale. Metabolites with blank circles were not resolved by the metabolomics. (**I**) Metabolite ratios mapped to the NAD^+^ biosynthesis pathway.

Metabolomics analysis yielded surprisingly few differentially expressed metabolites at baseline in these two mouse models (**Fig 3E**). Specifically, only enterolactone sulfate and chiro-inositol were elevated in the kidney of A-to-I locked mice compared to A-to-I uneditable mice. While the function of enterolactone sulfate remains unclear (weakly estrogenic (*43*)), chiro-inositol is a well characterized metabolite known to facilitate the conversion of pyruvate to acetyl-coA through the dephosphorylation of pyruvate dehydrogenase (*44, 45*). Dephosphorylation of pyruvate dehydrogenase is central to providing metabolic flexibility (*46*). Thus, the heightened chiro-inositol level could potentially explain the resilience of the A-to-I locked state against mitochondrial insults such as ischemia-reperfusion injury and direct ATP synthase inhibition as shown above (**Fig 2E**).

In contrast to basal conditions, we identified multiple differentially expressed metabolites following ischemia-reperfusion injury in these two mouse strains (**Fig 3F, G**). First, A-to-I locked state resulted in global upregulation of metabolites involved in the polyamine pathway, including S−adenosylmethionine, which serves as a donor of amine groups essential for the synthesis of higher order polyamines (spermidine and spermine; **Fig 3H**). Interestingly, A-to-I locked mice showed increased NAD^+^ levels following ischemia-reperfusion injury (**Fig 3I**). The beneficial effects of NAD^+^ have been extensively characterized across various animal models and human studies (*47*). We also found that A-to-I locked mice had higher levels of AICAR (5-Aminomidazole-4-carboxamide ribonucleotide), which originates from the pentose phosphate shunt/purine metabolism (**Fig 3F and Suppl Fig 4B**). The elevated AICAR levels under ischemic stress could result from the augmented glycolytic capacity conferred by the A-to-I locked condition. AICAR operates as a potent endogenous AMPK activator, contributing to a multitude of cellular protection mechanisms (*48*). Altogether, our findings indicate that Azin1 A-to-I editing renders cells resilient to ischemic stress by harnessing multiple protective pathways. These pathways encompass the upregulation of polyamine biosynthesis, NAD^+^ biosynthesis, and pentose phosphate shunt/purine metabolism. Finally, we examined the role of Azin1 A-to-I editing in the endotoxemia model and confirmed the renoprotective effects of A-to-I locked state (**Suppl Fig 4C-E**).

### Origin of double-stranded RNA species

A-to-I editing is catalyzed by an enzyme Adenosine Deaminase, RNA Specific (ADAR), which specifically binds to double-stranded RNA structures (dsRNA) (*49*). To investigate the nature of dsRNA species involved, we examined kidneys from our murine model of endotoxemia. Immunoblotting revealed an acute increase in dsRNA levels one hour after endotoxin challenge (**Fig 4A**). dsRNA may arise from repetitive elements resembling virus-like structures, such as long terminal repeats (LTR) and non-LTR retrotransposons (SINEs and LINEs) (*50*). PCR analysis of select repeat elements revealed an increase of MusD (type D murine LTR retrotransposons) 4 hours after endotoxin challenge in the kidney (**Suppl Fig 5A**). However, in general, our select PCR targets did not show consistent results, suggesting that the origin of dsRNA may not be repeat class specific.

**Figure 4.**
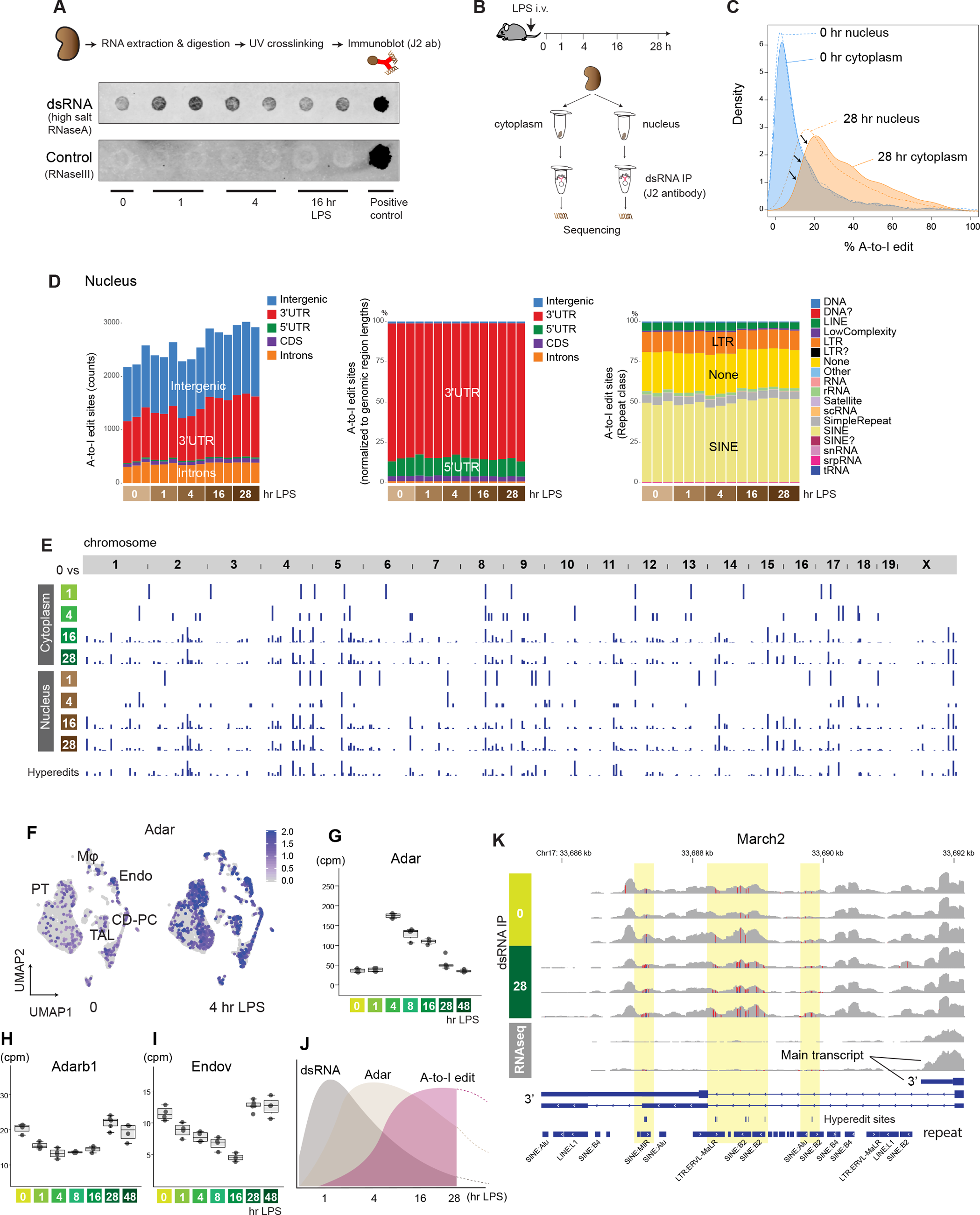
Genome-wide characterization of A-to-I editing in mouse kidneys. (**A**) Immunoblotting of dsRNA under the indicated conditions. RNaseA incubation was done with high salt to specifically digest single-stranded RNA. The negative control consisted of RnaseIII digestion, which digests dsRNA. The positive control consisted of poly(I:C) without RNase digestion. (**B**) Schematic representation of dsRNA immunoprecipitation and sequencing. (**C**) Overlay of density plots displaying A-to-I edit percentages under the indicated conditions. (**D**) Left: Total counts and distribution of A-to-I editing sites per sample (nuclear fraction; editing rate > 10% and reads count > 5 in at least 3 samples; refer to Methods for further preprocessing criteria). Middle: Distribution of A-to-I editing sites, normalized to genomic region lengths. Right: Distribution of A-to-I editing sites per repeat class. (**E**) Summary of A-to-I editing sites that exhibit differential expression compared to the 0-hour baseline. The bottom track represents hyper-editing sites. (**F**) Single-cell UMAP displaying the distribution of Adar expression in the mouse kidney (Reanalysis of published data GEO GSE151658). (**G**-**I**) Quantitation of total RNA-seq read counts (in counts per million) at the specified time points. (**J**) Scheme depicting the sequence of events observed in the kidney. (**K**) Read coverage comparison for March2 near the transcription termination site between dsRNA enrichment (top 6 tracks) and without dsRNA enrichment (bottom 2 tracks, 0 and 28 hours post LPS; regular total RNA sequencing).

To further investigate the phenomenon of dsRNA stress, we next conducted immunoprecipitation of dsRNA species, followed by stranded RNA-seq on cytoplasmic and nuclear fractions (dsRNA-seq; **Fig 4B, Suppl Fig 5B - F**) (*49*). The dsRNA-specific monoclonal antibody (J2) effectively enriched transcripts of varying lengths, ranging from ∼40 base pairs to several thousand base pairs. This is in line with the antibody’s known characteristics (*51*). In comparison to conventional total RNA-seq, dsRNA-seq enriched mitochondrially-encoded RNA transcripts at baseline and during the early stages of endotoxemia (**Suppl Fig 5E**). This observation is consistent with the fact that 1) mitochondrial transcription is highly active in the kidney at early time points (**Suppl Fig 5G**; mitochondrial transcription decreases at later time points), and 2) mitochondrial transcripts are prone to forming dsRNA structures due to the bidirectional transcription of the mitochondrial genome (*52*). Immunoprecipitation of dsRNA species also led to the enrichment of intergenic transcripts (**Suppl Fig 5E**). A more detailed examination revealed that these intergenic dsRNAs were particularly common in regions adjacent to the gene coding regions (±10 kb from transcription start and end sites; **Suppl Fig 5H**). These regions are prone to various mechanisms that can induce the generation of antisense reads, thereby facilitating dsRNA formation (*53, 54*).

In addition to antisense reads, intramolecular base pairing of single-stranded RNA (stem-loop) is another important source of dsRNA structures that can be catalyzed by ADAR. As demonstrated in **Supplemental Figure 5I**, sufficiently long complementary repeat regions are present in many genes, especially within intronic regions (912 genes for 30 bp cutoff). These repeat regions contribute to the enrichment of various gene body regions, including introns within our dsRNA-seq data set.

### Characterization of A-to-I editing sites

Having identified dsRNA species that could be targets of ADAR, we next examined the broad distribution of A-to-I editing sites in the mouse kidney. Our data revealed millions of A-to-I editing sites distributed across the genome (see Methods). However, the majority of these editing sites had low coverage (≤5 reads), minimal editing levels (a few percent), or inconsistent editing patterns per condition. Thus, we implemented stringent filtering criteria and focused our analysis on approximately 3,000 editing sites of high confidence for the rest of this study. Importantly, our analytical pipeline employed sequential alignment procedures (**Suppl Fig 5B**), enabling the capture of hyper-editing sites that will otherwise fail to map to a reference genome due to an excess of mismatches (*55*).

Across the genome, we found that both the extent of editing per site and the number of edited sites increased during the later stages of endotoxemia (**Fig 4C-E, Suppl Fig 6A, B**). A comparison of the cytoplasmic and nuclear fractions revealed that A-to-I editing occurred predominantly in the nucleus. Nevertheless, although the nucleus remained the primary site of editing, a greater number of transcripts underwent editing within the cytoplasm during the late phase of endotoxemia (**Fig 4C**). This transition was preceded by an upregulation of Adar expression across all cell types in the kidney (**Fig 4F-H**; the paralog of Adar, Adarb1, was downregulated). In parallel, the expression of Endonuclease V, the inosine-specific endoribonuclease, decreased, which would also contribute to the preservation of A-to-I edited transcripts (**Fig 4I**).

In summary, our comprehensive time-course analysis delineated the sequence of events leading to A-to-I editing: initiation with dsRNA stress (1 hr), followed by Adar overexpression (4 hrs), and culminating in an increase in A-to-I editing (16 hrs; **Fig 4J**). Representative dsRNA-seq read coverage tracks for each time point are available on a genome browser at https://connect.posit.iu.edu/view_GY/.

Over the entire endotoxemia time course, A-to-I editing was most prominent in 3’ untranslated regions (3’UTR; **Fig 4D, Suppl Fig 6A**). This prevalence of A-to-I editing in the 3’UTR was more pronounced in hyper-editing sites (**Suppl Fig 7A, B**). As expected, editing occurred preferentially in repeat regions, especially in SINE (**Fig 4D, Suppl Fig 6A, Suppl Fig 7B**). No significant temporal changes were observed in the overall proportion of edit sites per repeat class. We observed markedly different read coverage distribution between dsRNA-seq and total RNA-seq for certain genes. For example, March2, an E3 ubiquitin ligase involved in antiviral and antibacterial immune responses, showed significant transcription readthrough with respect to the canonical transcription termination site across a series of hyper-edited regions (**Fig 4J**). This phenomenon of readthrough was not readily apparent in conventional RNA-seq data, suggesting that these heavily edited transcripts might be lowly expressed or unstable (**Suppl Fig 7C, D**). The enrichment of dsRNA reads in intronic regions, specifically in repeat regions, was also notable (**Suppl Fig 7E, F**). The high prevalence of A-to-I editing in the intronic regions indicates that editing takes place immediately on nascent transcripts prior to splicing. Given that A-to-I editing in intronic regions could potentially impact alternative splicing (*56*), we further scrutinized individual editing sites. Nearly all edit sites were found outside of splicing donor or acceptor regions, including the branch point adenosine (**Suppl Fig7G**). Pathway enrichment analysis revealed that differentially edited sites are enriched in genes related to the regulation of ribonucleoproteins/P-bodies and the unfolded protein response/endoplasmic reticulum membrane (**Fig 5A, B**).

**Figure 5.**
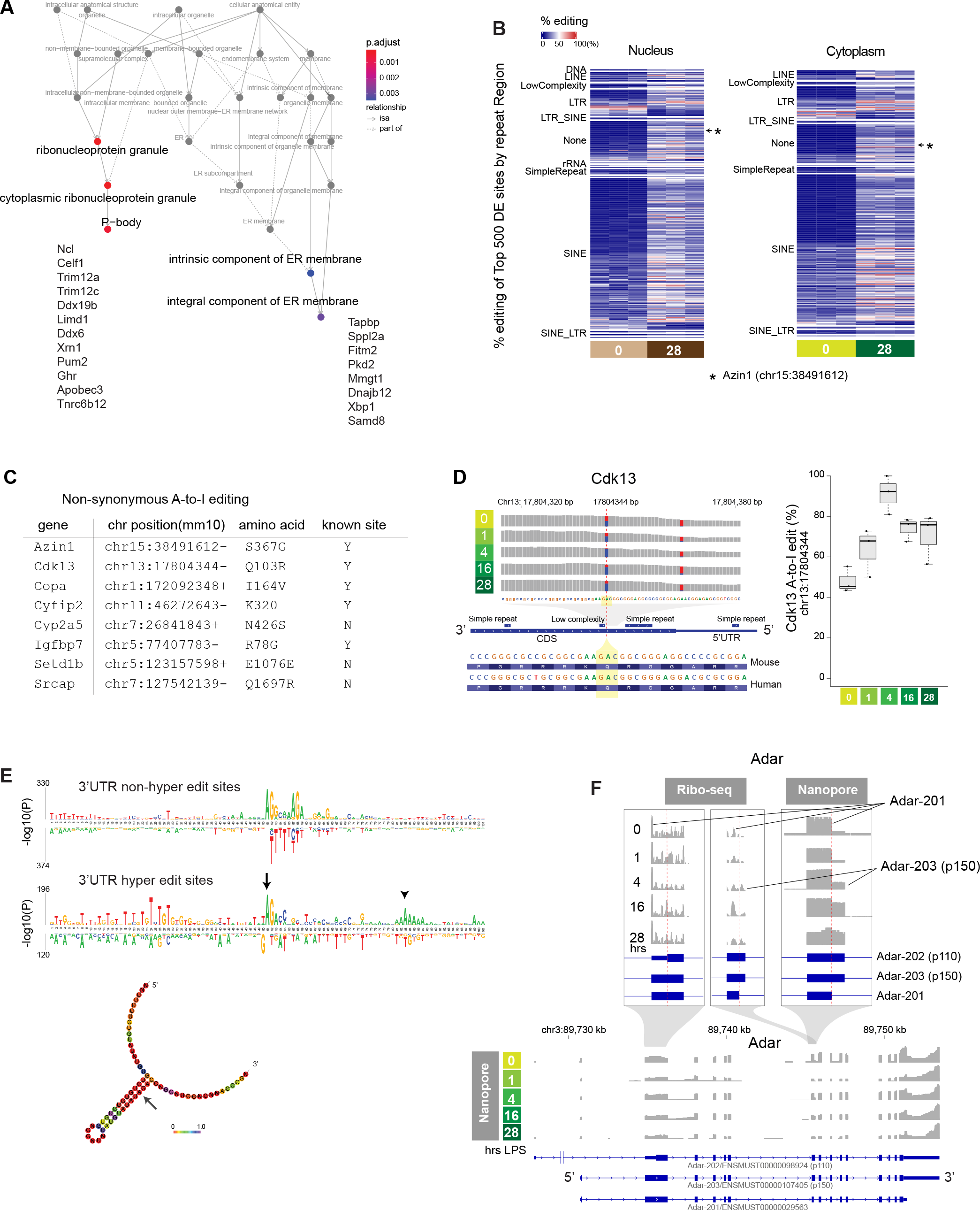
Genome-wide characterization of A-to-I editing in mouse kidneys. (**A**) Pathway enrichment analysis based on genes that exhibit differential editing rates between baseline and 28 hours (cytoplasmic compartment). (**B**) Heatmap displaying the top 500 differentially expressed A-to-I editing sites between 0-hour baseline and 28 hours after endotoxin in the kidney. The differentially expressed sites are categorized based on repeat classes. (**C**) List of genes exhibiting non-synonymous A-to-I coding sequence mutation in response to an endotoxin challenge in the kidney. (**D**) Cdk13 reads distribution and A-to-I editing under the indicated conditions. (**E**) Comparison of motif enrichment between non-hyper-editing (top) and hyper-editing sites (bottom) within ±50 nucleotides centered around A-to-I editing sites. Predicted RNA secondary structure around the 3’UTR hyper-editing site is shown at the bottom (arrow). (**F**) Ribo-seq and Nanopore read coverage graphs for Adar, clarifying Adar transcript isoform switches during endotoxemia.

### A-to-I editing within coding sequence regions

A-to-I editing within coding sequences was exceedingly rare. Specifically, instances of A-to-I editing that led to non-synonymous mutations were identified only in the following genes: Azin1, Cdk13, Copa, Cyfip2, Cyp2a5, Igfbp7, Setd1b, and Srcap (**Fig 5C**). Cdk13 functions as a transcriptional cyclin-dependent kinase involved in nuclear RNA surveillance. The A-to-I editing event in the Cdk13 coding sequence occurred near the N-terminus between two repeat regions, resulting in a glutamine-to-arginine mutation (**Fig 5D**). This particular editing site is conserved across both mice and humans (*34*). We found that the rate of editing at this site is markedly elevated at baseline and increased even further after endotoxin challenge (**Fig 5D**). Notably, this editing site was recently linked to aggressive cancer phenotypes (*57*), analogous to the findings with Azin1 A-to-I editing.

In the case of Azin1, editing at chromosome 15:38491612 (mm10) results in a serine-to-glycine mutation. The rate of Azin1 editing significantly increased from 0% to over 40% with the progression of endotoxemia (**Suppl Fig 8A, B**). When the serine-to-glycine mutation occurred, the neighboring adenosine was also edited in approximately 50% of cases, resulting in a synonymous mutation (chr15:38491613). Isolated editing of this adjacent adenosine was rare, confirming that chr15:38491612 is indeed the primary editing site. There was a complete absence of A-to-I editing two nucleotides away from the main editing locus. These findings underscore the remarkable precision and highly predictable nature of A-to-I editing. Because the Azin1 editing site is located near the alternative splice site, we also conducted Nanopore long-read RNA-sequencing and determined that the Azin1 editing status does not correlate with alternative splicing (**Suppl Fig 8C, D**).

### Adar isoform switching in the mouse kidney

While the clinical implications of A-to-I editing at individual sites remain largely unknown and some are likely inconsequential, various studies have underscored the significance of A-to-I editing in controlling the kinetics of transcripts, RNA-RNA interactions, R-loop formation, and RNA-protein interactions (*58, 59*). Generally, A-to-I editing serves to disrupt a long stretch of complementary base pairing, thereby attenuating the binding of dsRNA sensors such as PKR and MDA5 (*60*). Our motif enrichment analysis revealed ADAR’s preference for editing adenosines adjacent to guanosines (5’AG3’; **Fig 5E**), consistent with prior reports (*61, 62*). Intriguingly, when focusing on hyper-editing sites, we detected satellite A-rich regions situated approximately 30 nucleotides downstream of the primary editing site within the 3’UTR (**Fig 5E bottom**). ADAR has been shown to edit recursively at a fixed interval of approximately 30 base pairs downstream of an editing site (*63*).

Using Nanopore long-read RNA-sequencing and Ribo-seq, we identified that a relatively less characterized isoform Adar-201/ENSMUST00000029563 predominates in the kidney at baseline and up to 4 hours after endotoxin treatment (**Fig 5F**). The more widely recognized Adar isoforms are p110 (Adar-202/ENSMUST00000098924) and p150 (Adar-203/ENSMUST00000107405) (*64*). The constitutively expressed p110 (in other tissues) lacks a nuclear export signal, hence exerting its editing effect almost exclusively within the nucleus. In contrast, the interferon-inducible isoform p150 harbors both nuclear export and nuclear localization signals, enabling its shuttling between the cytoplasm and nucleus. Similarly, the 201 isoform possesses both nuclear export and nuclear localization signals identical to p150, permitting Adar-201 to distribute in both compartments. However, unlike the p110 and p150 isoforms, exon 7 of Adar-201 is truncated by 26 amino acids due to alternative splicing. This splicing occurs at the juncture of the critical dsRNA-binding domain, R_III_ (*65*). Therefore, it could potentially disrupt the editing capacity and account for the absence of significant editing by Adar in the kidney tissue at baseline.

In summary, our comprehensive analysis of endotoxemia model provided a timeline-specific landscape of A-to-I editing, represented by an array of novel and established editing loci. The phenomenon of A-to-I editing is highly reproducible and quantitative, thus offering significant potential for the development of a novel diagnostic and staging strategy for kidney disease.

## Discussion

The fast and variable progression of AKI poses a major challenge in implementing a stage-specific therapy at the bedside. We have previously identified that translation shutdown is a hallmark of late-phase septic AKI (*38, 40*). While transient inhibition of protein synthesis could be cytoprotective as it attenuates energy consumption and upregulates the integrated stress response, persistent inhibition of protein synthesis is detrimental. Importantly, in a reversible model of AKI, this late phase is also a crucial transition period where tissue recovery begins (*39*). How the tissue, under severe stress, reboots and attains a recovery phenotype is unclear.

In this study, we demonstrate that AZIN1 A-to-I editing plays a key role in promoting tissue recovery after AKI. Leading up to this robust AZIN1 editing is a series of stress responses the kidney goes through. These include NFκB-mediated acute inflammation, interferon responses and the integrated stress response, all culminating in metabolic shutdown (*38–40, 66*). Thus, AZIN1 A-to-I editing represents a landmark outcome following prolonged cellular stress. We found that the lack of AZIN1 editing renders cells susceptible to nutrient deprivation and attenuates glycolytic reserve, thereby restricting cell proliferation. Conversely, AZIN1 A-to-I editing confers better fitness by coupling increased polyamine bioavailability to the activation of cytoprotective molecules such as NAD^+^ and AICAR. The phenotypic impact of AZIN1 A-to-I editing *in vivo* is subtle under basal conditions but becomes apparent during stress. This indicates that AZIN1 A-to-I editing itself is not a driver of metabolic rewiring but assists this process during emergency. Collectively, these findings suggest a general model in which AZIN1 A-to-I editing serves as a rational auto-regulatory system, safeguarding against sustained metabolic shutdown and providing a cue for tissue recovery.

Our study also provides a comprehensive map of A-to-I editing in the kidney using a model of endotoxemia. This model is highly reproducible and has been extensively characterized (*14, 38, 39, 67–69*). This model was also independently benchmarked by Zhou et al. against a range of kidney injury models (*70*), further confirming the distinct stage transition from injury to recovery captured by this model. Our genome-wide interrogation of A-to-I editing revealed that A-to-I editing was enriched in genes involved in crucial stress response pathways including P-bodies and the unfolded protein response during the recovery phase of kidney injury (e.g., Limd1, Celf1, Pum2, Apobec3, Sppl2a, Dnajb12, and Xbp1). Given the biological relevance, these editing sites might have evolved to diversify their transcript repertoires or to evade the recognition by double-stranded RNA sensors under stress conditions. The latter mechanism has been clearly demonstrated for A-to-I editing in the kidney disease risk gene APOL1 by Riella et al (*71*).

Infections and various environmental factors frequently act as triggers and exacerbate the progression of kidney disease. The resulting outcomes exhibit significant variability. The present study portrays a timeline-specific role for A-to-I editing in the kidney during periods of stress. This structured transcriptional variation is quantitative and tied to an individual’s unique past. While not all the editing sites are necessarily pertinent or carry biological significance, it is our hope that further clarification of these attributes will enhance the accuracy of disease diagnosis and provide a molecular clock to guide therapy.

## Methods

### Malaria cohort

RNA-seq fastq files were obtained from GEO GSE52166 (stranded total RNA-seq with 2×100 bp paired-end configuration). The study details have been outlined previously (*33*). This longitudinal cohort study consisted of bi-weekly active malaria surveillance and passive surveillance through self-referral over a three-year period. RNA-sequencing and PCR were conducted on whole blood samples obtained from subjects before and after Plasmodium falciparum infection as determined through the prospective surveillance program.

### Human kidney biopsy

All ethical regulations were complied with as related to this study. Human sample experiments followed relevant guidelines and regulations. Bulk RNA-seq data files were obtained from 2 sources: the Biopsy Biobank Cohort of Indiana (GSE139061) and the Kidney Precision Medicine Project Atlas (https://atlas.kpmp.org/repository; accessed January 25, 2023) (*35, 36, 72, 73*). Tissues from the Biopsy Biobank Cohort of Indiana were acquired under waiver of informed consent. The Kidney Precision Medicine Project participants were consented. These bulk kidney tissues were process from an OCT block using the SMARTer Stranded Total RNA-Seq Kit v2 (Takara). Sequencing was performed in a 2×75 bp paired-end configuration using a NovaSeq platform (Illumina).

### Generation of AZIN1 A-to-I locked and A-to-I uneditable homozygous clonal cell lines

We designed single guide RNAs and single-stranded oligo DNA nucleotides (ssODN; homology directed repair donor oligos), and generated knock-in HEK293T cell lines using the CRISPR/Cas9 system. A target knock-in and protospacer adjacent motif block (PAM synonymous mutation; GTG/Valine to TGC/Valine) were introduced in the vicinity of the double-strand break (±10 bp) using the asymmetric donor DNA strategy (36 bp|cut|91 pb for the nontarget strand).

sgRNA (+PAM) used for both A-to-I locked and uneditable genome editing

5’TGATGAGCTTGATCAAATTG(TGG)3’

ssODN (antisense strand) for A-to-I locked state (AGC/Serine to GGC/Glycine):

5’GCAGATGGTTCATGGAAAGAATCTGCTCCCATGTTATCAAAGATAAGCCAATCTCCCACA TTCAGCTCAGGAAGAAGACAGC**c**TTC**g**ACAATTTGATCAAGCTCATCACAGGATGGACCCC AAAGGC3’

ssODN (antisense strand) for A-to-I uneditable state (AGC/Serine to TCC/Serine):

5’GCAGATGGTTCATGGAAAGAATCTGCTCCCATGTTATCAAAGATAAGCCAATCTCCCACA TTCAGCTCAGGAAGAAGACAG**ga**TTC**g**ACAATTTGATCAAGCTCATCACAGGATGGACCCC AAAGGC3’

Cells were cultured in 10 cm plates to 70% confluency prior to nucleofection. Approximately 150 x 10^3^ cells (5µL) were mixed with 1.49 µL of ssODN (100 µM) and Cas9 complex consisting of 18 µL SF 4D-nucleofector X solution + supplement1 (Lonza V4XC-2012), 6 µL of sgRNA (30 pmol/µL), and 1 µL of Cas9 2NLS nuclease, S. *pyogenes* (20 pmol/µL, Synthego). Nucleofection was done using Amaxa 4D-Nucleofector X (CM-130 program; Lonza). Cells were seeded in 15 cm plates at various concentrations. Clonal isolation was done manually. DNA extraction was done using Quick DNA Miniprep kit (Zymo Research, D3025). PCR was done using Q5 High-Fidelity DNA polymerase (NEB) and Monarch PCR Cleanup Kit (NEB, T1030). PCR primers used: 5’ ACTCACAAATTCAATACCTGCGT3’ (forward) and 5’TGCCTTAAAATAAAATCACCTTACCA3’ (reverse).

PCR products were electrophoresed in 2% agarose gel (TopVision Agarose Tablets ThermoFisher R2801) and bands were excised and extracted using QIAQuick Gel Extraction kit (Qiagen, #28706). Sanger sequencing was done at GeneWiz. Software used for design and analysis of mutant cell lines include CRISPRdirect, SnapGene, Primer3Plus, NEB Tm calculator, and Synthego ICE. Successful homozygous mutant cell lines were chosen for downstream experiments.

### Generation of Azin1 A-to-I locked and A-to-I uneditable mouse models

Similar to the human cell lines, we designed the following sgRNA and ssODN to generate A-to-I locked and A-to-I uneditable mouse models.

sgRNA (+PAM) used for both A-to-I locked and uneditable genome editing

5’TGATGAGCTTGATCAAATTG(TGG)3’

ssODN (antisense strand) for A-to-I locked state (AGC/Serine to GGC/Glycine):

5’GCAGATGGTTCGTGGAAAGAATCTGCTCCCATGTTATCAAAGATAAGCCAATCTCCCACA TTCAGCTCAGGAAGAAGACAGC**c**TTC**a**ACAATTTGATCAAGCTCATCACAGGATGGACCCC AAAGGC3’

ssODN (antisense strand) for A-to-I uneditable state (AGC/Serine to TCC/Serine):

5’GCAGATGGTTCGTGGAAAGAATCTGCTCCCATGTTATCAAAGATAAGCCAATCTCCCACA TTCAGCTCAGGAAGAAGACAG**ga**TTC**a**ACAATTTGATCAAGCTCATCACAGGATGGACCCC AAAGGC3’

Embryo manipulation and generation of founder mice on C57BL/6J background were performed by the Jackson Laboratory Mouse Model Generation Services. N1 sperms (heterozygous) have been cryopreserved at the Jackson Laboratory (Stock No. 414244 for Azin1 A-to-I locked and Stock No. 413737 for Azin1 A-to-I uneditable mice). PCR primers used for genotyping: 5’TGAGACTTATGCCTGATCGTTG3’ (forward) and 5’ GGTTCGTGGAAAGAATCTGC3’ (reverse).

### Animal models of kidney injury

Azin1 A-to-I locked and uneditable mice were housed at Indiana University School of Medicine under a 12 h light/dark cycle at 25 °C. For studies that did not require the knock-in mice, C57BL/6J mice were obtained from the Jackson Laboratory. All mice were 8–12 weeks of age and weighed 24–32 g. Male mice were used unless indicated. Animals were subjected to a single-dose, 4 mg/kg endotoxin (LPS) tail vein i.v. injection in a volume of 300 µl (*E*. *coli* serotype 0111:B4, Millipore Sigma). Untreated mice were administered an equivalent volume of sterile normal saline as a vehicle. Ischemia-reperfusion injury was performed under isoflurane anesthesia. Prior to surgery, mice were given extended-release buprenorphine at a dose of 3.25 mg/kg. The mice were subjected to a 20-minute bilateral renal pedicle clamp followed by reperfusion. A heating pad was used to ensure their rectal temperature remained above 36 °C throughout the surgical procedure. No antibiotics or fluid resuscitation were administered. Nicotinamide (400 mg/kg; Millipore Sigma) was injected intraperitoneally at the time of LPS injection.

### Cells

HEK293T cells, AZIN1 A-to-I locked and uneditable homozygous clonal cells were cultured in Dulbecco’s Modified Eagle Medium (4.5 g/L glucose, L-glutamine and Na pyruvate; Corning, 10-013-CV) with 10% fetal bovine serum (Midwest Scientific USDAFBS) and 100 U/mL penicillin and 100 µg/mL streptomycin (Thermo Fisher). All cell types were cultured at 37 °C with 5% CO_2_.

### dsRNA UV crosslinking and immunoblotting

Total RNA extraction was done on mouse kidney lysate using the RNeasy Plus Universal Midi kit (Qiagen). RNase digestion was then performed on the RNA extract using the following RNases. RNase A (Thermo Fisher EN0531) cleaves single-stranded RNA when NaCl is greater than 0.3 M. RNase A (1.5 µg in 1.5 µl) was added to 3.5 µL of 5M NaCl and 15 µg of RNA extract in a total volume of 50 µl and incubated at room temperature for 10 min. RNase III (Thermo Fisher AM2290) cleaves double-stranded RNA. RNase III (15 U in 15 µl) was mixed with 15 µg of RNA extract in 5 µl of 10X RNase III reaction buffer consisting of 500 mM NaCl, 100 mM Tris pH 7.9, 100 mM MgCl_2_, 10 mM DTT, brought up to a total volume of 50 µl, and incubated at 37 °C for 1 hour. For both RNase A and RNase III digestion, the reaction was deactivated by adding 300 µl of TRIzol. Digested RNA was purified using Direct-zol (Zymo Research), and then 20 units of Superase*In was added to each sample (Thermo Fisher) before preparing a serial dilution. As a positive control, double-stranded RNA poly(I:C) (Invivogen) was used. An equal volume (2.5 µl) of purified RNA was dotted on Amersham Hybond N+ membrane (VWR) and crosslinked using an UV Stratalinker 2400 (Autocrosslink mode). Odyssey Blocking buffer was used for blocking. The membrane was incubated with anti-dsRNA monoclonal antibody J2 (SCICONS/Jena Bioscience RNT-SCI-10010200) at a concentration of 2.5 µg/mL overnight at 4 °C followed by anti-mouse secondary antibody.

### dsRNA immunoprecipitation and RNA-Seq

We adopted and made modifications to existing protocols (*52, 74*). Mouse kidneys were harvested and immediately minced on an ice-cold dish. One-third of the minced tissue was transferred to a 1.2 mL of fractionation/lysis buffer, which consisted of 10 mM Tris pH 7.0, 10 mM NaCl, 5 mM MgCl_2_, 0.5% IGEPAL CA-630 (Sigma I8896), 0.5% Triton X-100, DNase I (Zymo E1011A 10 U/mL), and Superase*In (2 uL per 1 mL; Thermo Fisher). The lysate was centrifuged at 3,000 g for 3 minutes at 4 °C. The supernatant was further centrifuged at 21,000 g for 5 minutes. The resulting supernatant represents the cytoplasmic fraction. The pellet obtained from the initial centrifugation was resuspended in 1 mL of the fractionation/lysis buffer and homogenized using a Minilys tissue homogenizer (Bertin Instruments) at the highest speed for 45 seconds. After homogenization, the lysate was centrifuged at 21,000 g for 5 minutes. This supernatant serves as the nuclear fraction. Each fraction was then incubated with anti-dsRNA monoclonal antibody J2 (SCICONS/Jena Bioscience RNT-SCI-10010200; IgG2a kappa light chain) at a concentration of 10 µg per 600 µl of lysate for 2 hours at 4 °C. Mouse IgG2a kappa (clone eBM2a; eBioscience 14-4724-82) was used as an isotype control. Protein G Dynabeads (Invtrogen, 10003D) were washed in the immunoprecipitation buffer described below and then incubated with the sample-antibody mix for 1 hour at 4 °C. The dsRNA-antibody-Dynabeads complex was washed on a magnetic rack using 500 µL x 4 immunoprecipitation buffer consisting of 50 mM Tris pH 7.4, 100 mM NaCl, 3 mM MgCl_2_, IGEPAL 0.5%. RNA was extracted from Dynabeads using 1 mL TRIzol and 200 µL chloroform per sample. After the second round of TRIzol chloroform RNA purification, RNA precipitation was done using ice cold isopropanol, sodium acetate and GlycoBlue on ice for 1 hour. The RNA was resuspended in 7 µL of water. The RNA yields were approximately 6 ng/µl to 16 ng/µl for J2 antibody immunoprecipitation (lower in the nuclear fraction), while the isotype control yielded less than 300 pg/µl.

Stranded total RNA-sequencing was performed at the Indiana University Center for Medical Genomics Core. cDNA library preparation was carried out using the Clontech SMARTer Stranded Total RNA-Seq Kit v2. The pooled libraries were loaded onto a NovaSeq 6000 sequencer at 300 pM final concentration for 100 bp paired-end sequencing (Illumina). Approximately 40 million reads per library was generated. Phred quality score (Q score) was used to measure the quality of sequencing. More than 90% of the sequencing reads reached Q30 (99.9% base call accuracy). The sequenced data were mapped to the mm10 or hg38 genome using STAR. The counts data were generated using featureCounts and analyzed using edgeR (https://github.com/hato-lab/A-to-I-edit).

### A-to-I editing analysis

The entire data processing scripts are available through GitHub: https://github.com/hato-lab/A-to-I-edit.

Fastq files were initially aligned to the reference genomes: GRCm38 primary assembly and Gencode vM25 gtf files for mouse and GRCh38 and v41 gtf files for human, using the STAR aligner (v2.7.9a).

STAR --runMode alignReads --outSAMtype BAM SortedByCoordinate --readFilesType Fastx -- readFilesIn $first_antisense_mates $second_sense_mates --readFilesCommand zcat -- limitBAMsortRAM $params.mem_bytes --outFileNamePrefix $prefix --outReadsUnmapped Fastx --outMultimapperOrder Random --outBAMsortingThreadN $threads

To capture hyper-edited reads (*55*), we generated pseudo-genome references where all ‘A’ bases were substituted with ‘G’ (see **Supplemental Figure 5A**). The unaligned reads from the initial alignment were realigned to the pseudo-genome reference using the STAR aligner with the same parameters (hyper-edited reads).

A-to-I editing sites were first detected using reditools2 extract_coverage.sh and parallel_reditools.py (*75*). The resulting A-to-I editing sites underwent additional filtering based on the following criteria: for a given editing site, there must be at least 3 samples with edited reads greater than 5 and an editing rate greater than 0.1 but less than 0.9 in order to reduce the inclusion of low editing loci and potential genomic mutations, respectively. The reading depth of the remaining sites (total counts) was obtained using the samtools depth command with the -b option (v1.9). Each A-to-I editing site was annotated using biomaRt and a repeat class file obtained from the UCSC Genome Browser. Fisher’s exact test was employed to compare edit ratio group comparisons. P-values for each comparison were adjusted using the false discovery rate method. Sites with false discovery rate adjusted p-values <0.05 were considered significant for each comparison. Annovar was utilized to identify coding sequence mutations, and motif enrichment analysis was done using kpLogo.

Complementary repeat region analysis was done using Biostrings::findPalindromes in R. RNAfold was used for the prediction of RNA minimum free energy secondary structures.

### Nanopore long-read RNA sequencing

Snap-frozen mouse kidney tissue was homogenized in 800 µl of Tri Reagent using a Minilys tissue homogenizer at the highest speed for 45 seconds and then incubated for 5 minutes. Total RNA was extracted from 600 µl of the supernatant using the Direct-zol RNA miniprep Plus (Zymo Research), including on-column DNase I digestion. The RNA was eluted in 100 µl of water and subjected to mRNA polyA enrichment using the Dynabeads mRNA DIRECT Micro kit (Thermo Fisher 61021). After two rounds of washing and enrichment of mRNA following the manufacture’s protocol, polyA+ mRNA was eluted in 10 µl of water.

To establish our Nanopore RNA sequencing workflow, we initially compared direct RNA sequencing (SQK-RNA002), direct cDNA sequencing (SQK-DCS109), and cDNA-PCR sequencing (SQK-PCS111), all on R9.4.1 flow cells. We chose direct cDNA sequencing because: 1) direct RNA sequencing had lowest read coverage and 3’ reads bias, and 2) cDNA-PCR sequencing had shortest reads length (but the highest reading depth). We proceeded with approximately 200 ng of polyA+ mRNA as the starting material. Reverse transcription and strand-switching was done using Maxima H Minus Reverse Transcriptase following the Nanopore protocol (SQK-DCS109). After the second strand synthesis, the RNA concentration was generally 20-30 ng/µl as measured by NanoDrop. Following end-prep, adapter ligation, and AMPure XP bead binding, the yield was generally 6-10 ng/µl as measured by Qubit. Sequencing was conducted on R9.4.1 flow cells using the GridIon platform.

The basecalling was done using Guppy basecaller:

guppy_basecaller --compress_fastq --fast5_out -i ./fast5_pass/ -s ./fastq/ --device ‘auto’ -- num_callers 1 --flowcell “FLO-MIN106” --kit “SQK-DCS109”

Mapping was carried out using Minimap2

minimap2 -a -L -t 12 -x splice --junc-bed Mus_musculus.GRCm38.101.bed --MD mm10-ont.mmi ./fastq/*.fastq.gz 2> ./minimap2.err > ./SAM/$sample.sam

Subsequent transcript-level analysis was performed using Bambu (*76*).

### Transformation of E. coli and plasmid transfection of HEK293T cells

We designed the following two plasmids based on the pCDH-EF1a-eFFly-mCherry backbone vector (Addgene #104833) with cloning sites BmtI and BamHI. Gene synthesis was done by GenScript, and the resulting plasmid sequences were verified using Sanger sequencing.

pCDH-EF1a-3xFLAG-AZIN1_locked-T2A-mCherry

pCDH-EF1a-3xFLAG-AZIN1_uneditable-T2A-mCherry

The sequence of 3xFLAG tag consisted of: gccaccatg(Kozak sequence)gactacaaagaccatgacggtgattataaagatcatgacatcgattacaaggatgacgatgacaagAAA(AZIN1 CDS in frame).

The sequences of AZIN1-locked and uneditable plasmids differ at the A-to-I editing site, following the same mutation strategy used in the CRISPR knock-in cell lines.

A-to-I locked: 5’TGATGAGCTTGATCAAATTGT**c**GAA**g**GC3’

A-to-I uneditable: 5’TGATGAGCTTGATCAAATTGT**c**GAA**tc**C3’

The stop codon of AZIN1 is removed and fused in frame with T2A and mCherry CDS as follows: GCT(end of AZIN CDS without a stop codon)ggatccgcggccgctgagggcagaggaagtcttctaacatgcggtgacgtggaggagaatcccggccctatgcatATG(mCherry CDS start).

Chemically competent E. coli (One Shot Top10, Invitrogen C404003) were transformed with pCDH plasmids using the heat shock method (on ice for 30 min followed by 42 °C heat shock for 45 seconds). Transformed E. coli were incubated overnight in LB broth with 100 µg/mL carbenicillin at 225 rpm, 37 °C. Plasmid isolation was done using Nucleobond Xtra Midi Kit (Takara #740410.50) following the manufacture’s protocol. For DNA gel electrophoresis, 2% agarose gel was prepared using TopVision Agarose Tablets (Thermo Fisher R2801) and ethidium bromide (Thermo Fisher 15585011). Samples were electrophoresed with NEB X6 gel loading dye in 0.5X TAE buffer with a TrackIt 100 bp DNA ladder at 100 V. Transfection of plasmids was done using lipofectamine 2000 following the manufacturer’s instruction (sequential mixing of lipofectamine, Opti-MEM and plasmid into freshly replaced DMEM/10% FBS medium, Thermo Fisher 11668027). For all experiments, either 4 µg or 20 µg plasmid DNA was used per well of a 6 well-plate or 10 cm dish, respectively.

### AZIN1 immunoprecipitation and mass spectrometry

Cells were cultured in 10-cm plates to 80% confluency and lysed using 1 ml of 1X dilution Cell Signaling Cell lysis buffer (#9803), which consists of 2 mM Tris-HCl (pH 7.5), 15 mM NaCl, 0.1 mM Na_2_EDTA, 0.1 mM EGTA, 0.1% Triton, 0.25 mM sodium pyrophosphate, 0.1 mM beta-glycerophosphate, 0.1 mM Na_3_VO_4_, 0.1 µg/ml leupeptin, 1 mM PMSF (Thermo Fisher 36978), phosStop (1 pill per 10 ml), and protease inhibitor (1 pill per 10 ml; cOmplete Mini EDTA-free, Roche Diagnostics). The lysates were sonicated and then centrifuged at 14,000 x *g* for 10 minutes at 4°C. In a separate experiment, nuclear and cytoplasmic fractionation was done using 1 mL Pierce IP Lysis Buffer (87788). The cell suspension was passed through a 27-gauge needle 10 times using a 1mL syringe and incubated on ice for 20 minutes. The lysed cell suspension was then centrifuged at 720 x g at 4°C for 5 minutes. The resulting supernatant was collected as the cytoplasm fraction. The pellet was resuspended with 1 mL Pierce IP Lysis Buffer and passed through a 25-gauge needle 10 times, followed by centrifugation at 21,000 x g at 4°C for 15 min. The resulting supernatant was collected as the nuclear fraction.

FLAG-tag immunoprecipitation was performed by incubating 900 µL of the supernatant with 50 μL of pre-washed Pierce DYKDDDDK Magnetic Agarose (12.5 μL settled magnetic agarose; Thermo Fisher A36797) at room temperature for 20 minutes. The beads were washed three times with 1mL of phosphate buffered saline followed by urea denaturation, TCEP reduction, CAA alkylation, and trypsin/LysC digestion (Promega V5072). After digestion, samples were quenched with 2% formic acid. Peptides were separated on an Ultimate 3000 HPLC with loading on a 5 cm C18 trap column Acclaim PepMap 100 (3 µm particle size, 75 µm diameter; Thermo Scientific, 164946) followed by a 15 cm PepMap RSLC C18 EASY-Spray column (Thermo Scientific, ES900) and analyzed using a Q-Exactive Plus mass spectrometer (Thermo Fisher) operated in positive ion mode. Solvent B was increased from 5%-35% over 75 min, to 90% over 2 min, back to 3% over 2 minutes (Solvent A: 95% water, 5% acetonitrile, 0.1% formic acid; Solvent B: 100% acetonitrile, 0.1% formic acid). A data dependent top 15 method was used with MS scan range of 200-2000 m/z, resolution of 70,000, AGC target 3e6, maximum IT of 100 ms. MS2 resolution of 17,500, fixed first mass 100 m/z, normalized collision energy of 30, isolation window of 4 m/z, target AGC of 1e5, and maximum IT of 50 ms. Dynamic exclusion of 30 sec, charge exclusion of 1, 7, 8, >8 and isotopic exclusion parameters were used. Data were analyzed by Proteome Discoverer 2.5 and SEQUEST HT. A *Homo sapiens* reviewed protein database was downloaded from the Universal Protein Knowledgebase / Translated European Molecular Biology Laboratory (051322 and supplemented with frequently observed contaminants for a full tryptic search with a maximum of 3 missed cleavages). Precursor mass tolerance was set to 10 ppm and fragment mass tolerance set at 0.02 Da. Dynamic peptide modifications were oxidation on methionine, phosphorylation on serine, threonine, or tyrosine; acetylation, met-loss, and met-loss plus acetylation at protein N-terminus. Static modifications were carbamidomethylation on cysteines. Percolator false discovery rate filtration of 1% was applied to both the peptide-spectrum match and protein levels. Search results were loaded into Scaffold 5 for visualization and analysis.

### Polyribosomal profiling

For polyribosome profiling of tissues, cardiac perfusion was performed with 6 mL of cycloheximide (100 μg/ml in PBS, Sigma). Harvested tissues were immediately placed in a lysis buffer consisting of 1% Triton X-100, 0.1% deoxycholate, 20 mM Tris-HCl, 100 mM NaCl, 10 mM MgCl_2_, EDTA-free Protease Inhibitor Cocktail Tablet (Roche) and 100 μg/ml cycloheximide. Tissues were homogenized using a Minilys tissue homogenizer (Bertin Instruments). Tissue homogenates were incubated on ice for 20 minutes, then centrifuged at 9,600 *g* for 10 minutes. The supernatant was added to the top of a sucrose gradient generated by BioComp Gradient Master (10% sucrose on top of 50% sucrose in 20 mM Tris-HCl, 100 mM NaCl, 5 mM MgCl_2_, and 100 μg/ml cycloheximide) and centrifuged at 283,800 *g* for 2 hours at 4 °C. The gradients were harvested from the top in a Biocomp harvester (Biocomp Instruments), and the RNA content of eluted ribosomal fractions was continuously monitored with UV absorbance at 254 nm. For polyribosome profiling of cultured cells, cells were rinsed with cold PBS first, then cells were scraped using the lysis buffer on ice. The remaining procedure was identical to tissue polyribosome profiling.

### Metabolomics

Untargeted global metabolomic analysis was performed at Metabolon Inc. Snap-frozen mouse kidney tissues were processed following the Metabolon standard extraction method (60% methanol) and their UPLC-MS/MS pipeline. Targeted metabolomic analysis was performed at Creative Proteomics. Snap-frozen mouse kidney tissues were homogenized in 70% acetonitrile and the supernatant (50 µl) was incubated for 30 minutes at 30 °C with 200 µl of a solution consisting of 100 µl of 20 mM dansyl chloride for polyamine derivatization, 50 µl of dopamine-d3 as internal standard, and 50 µl of a pH 9 buffer. Analysis was done on an UPLC-MS/MS instrument with the positive-ion detection mode and with 0.1% formic acid in water (A) and acetonitrile/isopropanol (B) as the mobile phase for gradient elution (20% to 100% B in 15 minutes) at 0.3 ml/min and 50 °C. HPLC NAD^+^ measurements were conducted in our laboratory using an Agilent 1100 series system equipped with a UV detector set at 254 nm and a Sigma SPELCOSIL LC-18-T column (15cm x 4.6 mm with a particle size of 3 µm). Tissues were homogenized using 10% HClO_4_ (2 mL of 10% HClO_4_ per 0.2 grams tissue). After centrifugation, the supernatant was neutralized with one-third volume of 3 M K_2_CO_3_ and incubated on ice for 10 minutes to allow CO_2_ degassing with intermittent vortexing. Buffer (A) was composed of 0.05 M potassium phosphate buffer (38.5 ml of 0.5 M KH_2_PO_4_ and 61.5 ml of K_2_HPO_4_) and Buffer (B) consisted of 100% methanol (ramped from 0% to 15% B in 11 minutes).

### Seahorse bioenergetics assays

First, Seahorse XFp cell energy phenotype test (Agilent) was performed. The stressor mix recipe consisted of FCCP and oligomycin, each at a well concentration of 1 µM. After confirming that HEK293T cells, including AZIN1 A-to-I locked and uneditable mutants, are glycolytic, we proceeded to the XFp glycolysis stress test. Cells were seeded 24 hours prior to the stress test (8,000 cells in 100 µl of DMEM per well). A glucose-free assay medium was prepared by mixing 10 ml of Seahorse XF base medium minimal DMEM (Agilent 103334-100), 100 µl Na pyruvate (Sigma S8636), and 200 µl L-glutamine (gibco 25030-081), and the pH was adjusted to 7.4 using NaOH. The medium was replaced with the glucose-free assay medium following the Seahorse protocol. Port A consisted of glucose (final well concentration 10 mM), Port B contained oligomycin (2 µM), and Port C contained 2-deoxyglucose (50 mM).

### Real-time cell growth monitoring

Cells were seeded on the first day of live cell imaging at a density of 2,500 cells per well or 5,000 cells per well in 100 µl of DMEM on a 96-well plate. The Satorius IncuCyte system was used with either a 4X or 10X objective lens, and 2- or 4-hour imaging intervals for 5 days. In some experiments, urea (Sigma U5378) was added at a well concentration of 60 mM. DMEM SILAC medium (Thermo Fisher 88364) and DMEM without glutamine (Thermo Fisher 10313021) were used for indicated experiments. Cell growth quantitation was performed using the IncuCyte built-in masking algorithm.

### PCR

RNA was extracted using TRI Reagent and Direct-zol RNA MiniPrep Plus (Zymo Research R2070). Reverse transcription was done using High Capacity cDNA Reverse Transcription kit (ThermoFisher/Applied Biosystems 4368814). Conventional PCR was done using 1.8% agarose gel. PCR primers used include:

MuERV 5’TTTCTCAAGGCCCACCAATAGT3’ (forward) and 5’GACACCTTTTTTAACTATGCGAGCT3’ (reverse) (*77*)

MusD 5’GATTGGTGGAAGTTTAGCTAGCAT3’ (forward) and 5’TAGCATTCTCATAAGCCAATTGCAT3’ (reverse) (*78*)

Line1 5’TTTGGGACACAATGAAAGCA3’ (forward) and 5’CTGCCGTCTACTCCTCTTGG3’ (reverse) (*78*)

IAP 5’CTTGCCCTTAAAGGTCTAAAAGCA3’ (forward) and 5’GCGGTATAAGGTACAATTAAAAGATATGG3’ (reverse) (*78*)

### Western blotting

Proteins from cells and tissues were extracted using RIPA buffer (ThermoFisher Pierce) with 0.5M EGTA, 0.5M EDTA, DNase I (0.1 U/µL, Ambion AM2222), Halt protease inhibitors (Pierce), phosStop inhibitor (Roche), and benzonase nuclease (25 U/mL EMDMillipore 70746-4). Total protein levels were determined using a modified Lowry assay (Bio-Rad). Equal amounts of total proteins (10 - 20 μg) were mixed with NuPAGE LDS Sample Buffer (Thermo Fisher) containing 100 mM DTT and separated by electrophoreses on NuPage 4%–12% Bis-Tris gels, followed by transfer to PVDF membranes. Samples related to immunoprecipitation (input, flowthrough, and immunoprecipitated proteins) were lysed with Pierce IP Lysis Buffer (25 mM Tris-HCl pH 7.4, 150 mM NaCl, 1 mM EDTA, 1% NP-40, 5% glycerol). For AZIN1 protein turnover determination, cells were incubated with 20 µM MG132 for a total of 3 hours and 250 µg/ml cycloheximide for the indicated duration before cell lysis. Antibodies used include the following: Antizyme inhibitor 1 (Proteintech, rabbit polyclonal, 1:1,000 dilution, #11548-1-AP), Histone H3 (1B1B2, Cell Signaling, mouse monoclonal, 1:1,000 dilution, #14269), Alexa Fluor 680 AffiniPure donkey anti-rabbit and Alexa Fluor 790 AffiniPure donkey anti-mouse secondary antibodies (Jackson ImmunoResearch, 1:5,000 dilution #715-655-150, #711-625-152). Quantitation was done using Image Studio.

### Statistics

Data were analyzed for statistical significance and visualization with *R* software 4.1.0. Error bars show SD. For multiple comparisons, 1-way ANOVA followed by pairwise *t* tests was performed using the Benjamini and Hochberg method to adjust *P* values. All analyses were 2-sided, and a *P* value of less than 0.05 was considered significant.

### Data sharing

RNA sequencing data are deposited in the NCBI’s Gene Expression Omnibus database: dsRNA IP sequencing, GSE244941 (release status private. Representative read coverage tracks for the cytoplasmic fraction are available on a genome browser at https://connect.posit.iu.edu/view_GY/); Mouse kidney Nanopore PCR-free direct cDNA sequencing, GSE244942 (release status private); mouse kidney total RNA sequencing data (GSEXXXX, to be submitted). AZIN1 cell line RNA sequencing data are available at https://connect.posit.iu.edu/azin1/. Proteomics data will be deposited in ProteomeXchange (accession: xxxx, username xxxxxx password: xxxxx). Reanalysis of Ribo-seq and single-cell RNA-seq was performed using GSE120877 and GSE151658.

### Code availability

Scripts are available through GitHub: https://github.com/hato-lab/A-to-I-edit

### Study approval

All animal protocols were approved by the Indiana University Institutional Animal Care Committee and conform to the NIH Guide for the Care and Use of Laboratory Animals, National Academies Press. Human subjects work was approved by the Institutional Review Board of Indiana University School of Medicine (IRB 190657223).

## Acknowledgements

We thank Dr. Amber Mosley, Dr. Whitney Smith-Kinnamari, Dr. Chunna Guo, and Dr. Mandy Bittner at the Indiana University School of Medicine Proteomics Core, and Dr. Yunlong Liu, Dr. Hongyu Gao, Dr. Fang Fang, and Dr. Xialing Xuei at the Center for Medical Genomics, and former lab member Thomas W. McCarthy. Measurement of serum creatinine concentration was performed by Drs. John Moore, Yang Yan, et al. at the University of Alabama at Birmingham/UCSD O’Brien Center Core for Acute Kidney Injury Research (NIH P30DK079337) using isotope dilution LC-MS. This work was supported by NIH grants R01-AI148282 and Veterans Affairs Merit (BX002901) to TH, R01-DK107623 to PCD, U01DK114923 to PCD and MTE.

---

**Supplemental Figure 1.**
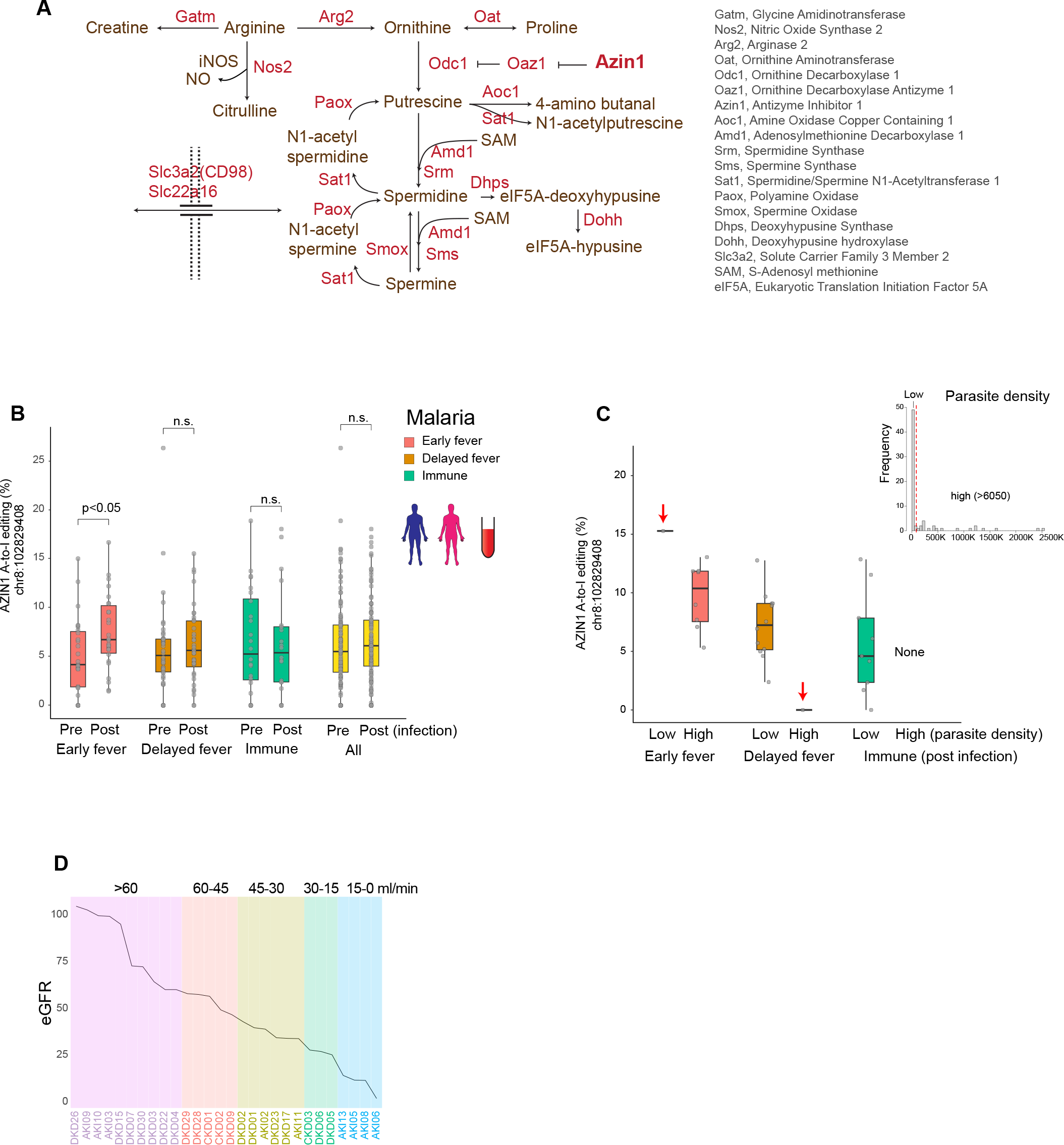
(**A**) Depiction of key molecules involved in the polyamine pathway. (**B**) Distribution of AZIN1 A-to-I editing rates (% of edited reads over total reads) for combined male and female individuals. (**C**) Distribution of AZIN1 A-to-I editing rates based on parasite density and clinical category for male children after malaria infection. The early fever group exhibited high parasite density, except for one child who had the highest AZIN1 editing rate and low parasite density. The delayed fever group had low parasite density, except for one child who had the lowest AZIN1 editing rate and high parasite density. All immune children had low parasite density. (**D**) Line plot of indicated conditions sorted by estimated glomerular filtration rate (eGFR). Detailed clinical data are not available, and the timing of kidney biopsy and the eGFR measurement is unclear.

**Supplemental Figure 2.**
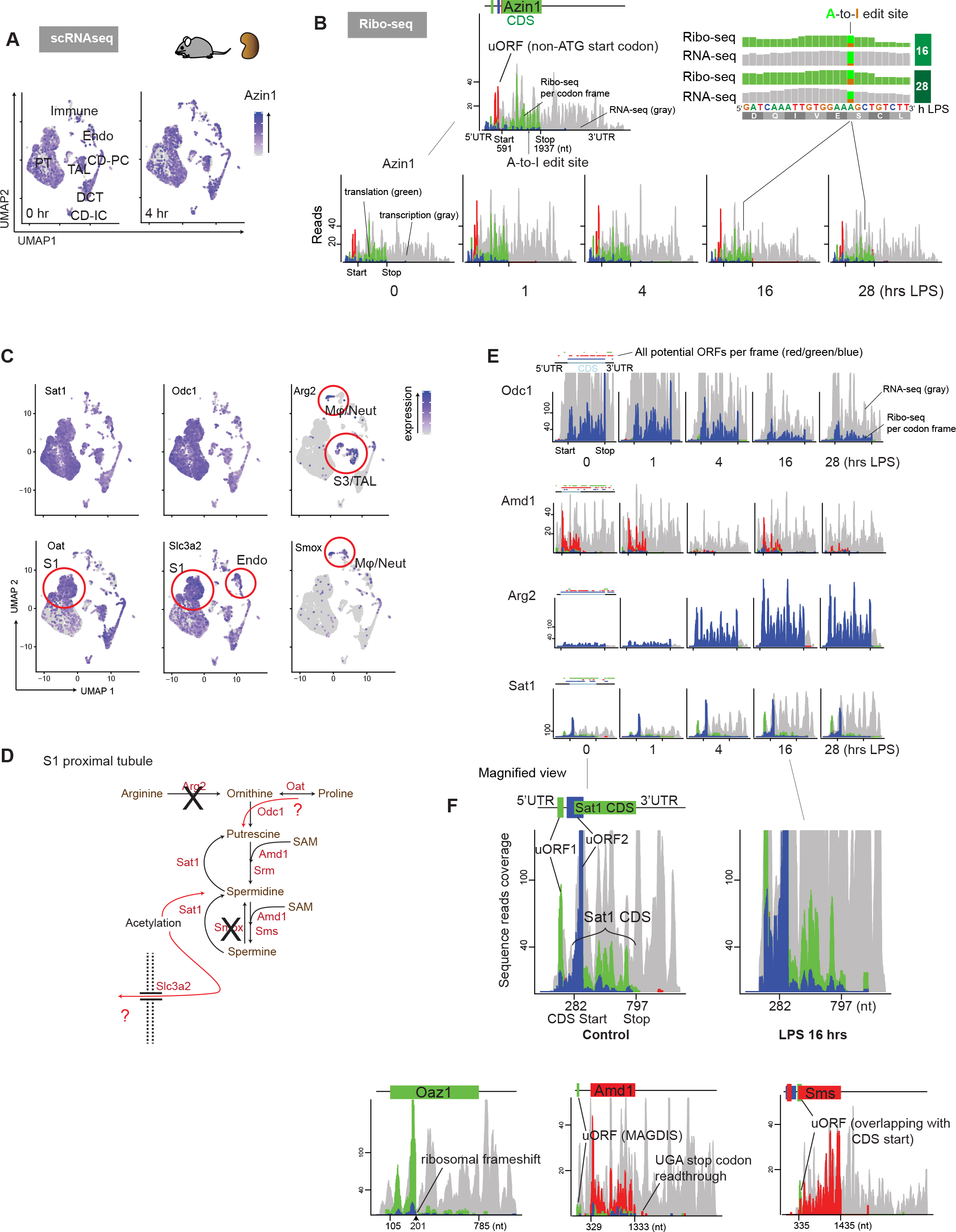
(**A**) Single-cell UMAP displaying the diffuse distribution of Azin1 expression in all cell types in the kidney (Reanalysis of published data GEO GSE151658). (**B**) Combined Ribo-seq and RNA-seq read coverage graphs for Azin1, showing the absence of significant changes in transcription and translation after endotoxin challenge in the kidney. Reads are mapped to ENSEMBL transcript Azin1-201. The gray-colored reads represent RNA-seq, whereas red/green/blue-colored reads represent codon frames for ribosome-protected fragments in Ribo-seq. The inset in the top right confirms the translation of A-to-I edited Azin1 (Reanalysis of published data GEO GSE120877). (**C**) Single-cell UMAP displaying gene expression levels and the distribution of indicated transcripts in the kidney. Select cell types are highlighted with red circles. (Smox and Oat are from 4 hours after LPS, and Arg2 is from 36 hours after LPS, the rest at from 0-hour baseline). (**D**) Interpretation of the single-cell RNA-seq data. S1 proximal tubules may preferentially utilize proline as a precursor for polyamines, whereas S3 proximal tubules rely more on arginine. (**E**) Combined Ribo-seq and RNA-seq read coverage graphs for the indicated genes (ENSEMBL Odc1-201, Amd1-201, Arg2-201, Sat1-201). Odc1 and Amd1 genes are polyamine anabolic enzymes, whereas Sat1 is a polyamine catabolic enzyme. (**F**) Annotation of distinct post-translational mechanisms in the polyamine pathway captured by Ribo-seq. uORF, upstream open reading frame; CDS, coding sequence.

**Supplemental Figure 3.**
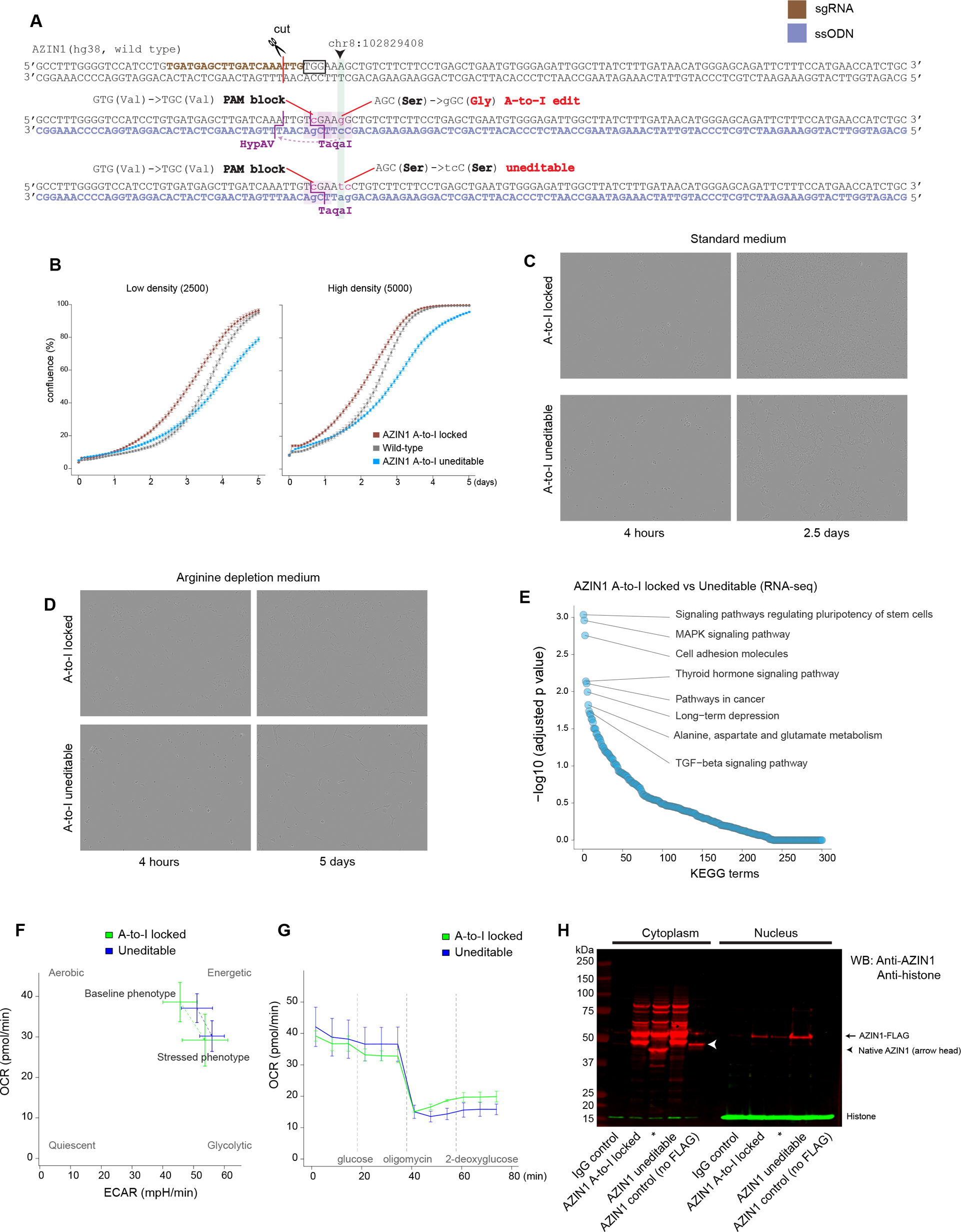
(**A**) CRISPR knock-in design used for generating the Azin1 A-to-I locked and uneditable cell lines. The single guide RNA is shown in brown, and single-stranded oligonucleotides are shown in blue. Key mutations introduced are annotated. (**B**) Comparison of cell growth between low-density and high-density seeding for AZIN1 A-to-I locked, uneditable, and wild-type cells (IncuCyte, 10x objective lens, imaged every 2 hours for 5 days. For clarity, high-density seeding data shown in the **Main** Figure 2B are presented again). (**C**, **D**) Representative time-lapse images under the indicated conditions. A-to-I uneditable cells exhibit delayed adherent cell morphology (compared at 4 hours post cell seeding). (**E**) KEGG pathway enrichment analysis comparing AZIN1 A-to-I locked and uneditable cell lines (RNA-seq data, n=3 technical replicates for each cell type). Approximately 300 KEGG metabolic pathway terms are aligned in the order of statistical significance. (**F**) Seahorse cell energy phenotype test. The x-axis denotes extracellular acidification rates, and y-axis denotes oxygen consumption rates. (**G**) Oxygen consumption rates corresponding to the **Main** Figure 2H Seahorse glycolysis stress test, confirming the effect of oligomycin in both cell lines. n=3 independent experiments with n=3 technical replicates for each experiment. (**H**) Western blotting for AZIN1 and histone, showing the expected shift of AZIN1 due to FLAGx3 as well as differential expression of histone between cytoplasmic and nuclear fractions. *denotes a plasmid construct not used in this manuscript.

**Supplemental Figure 4.**
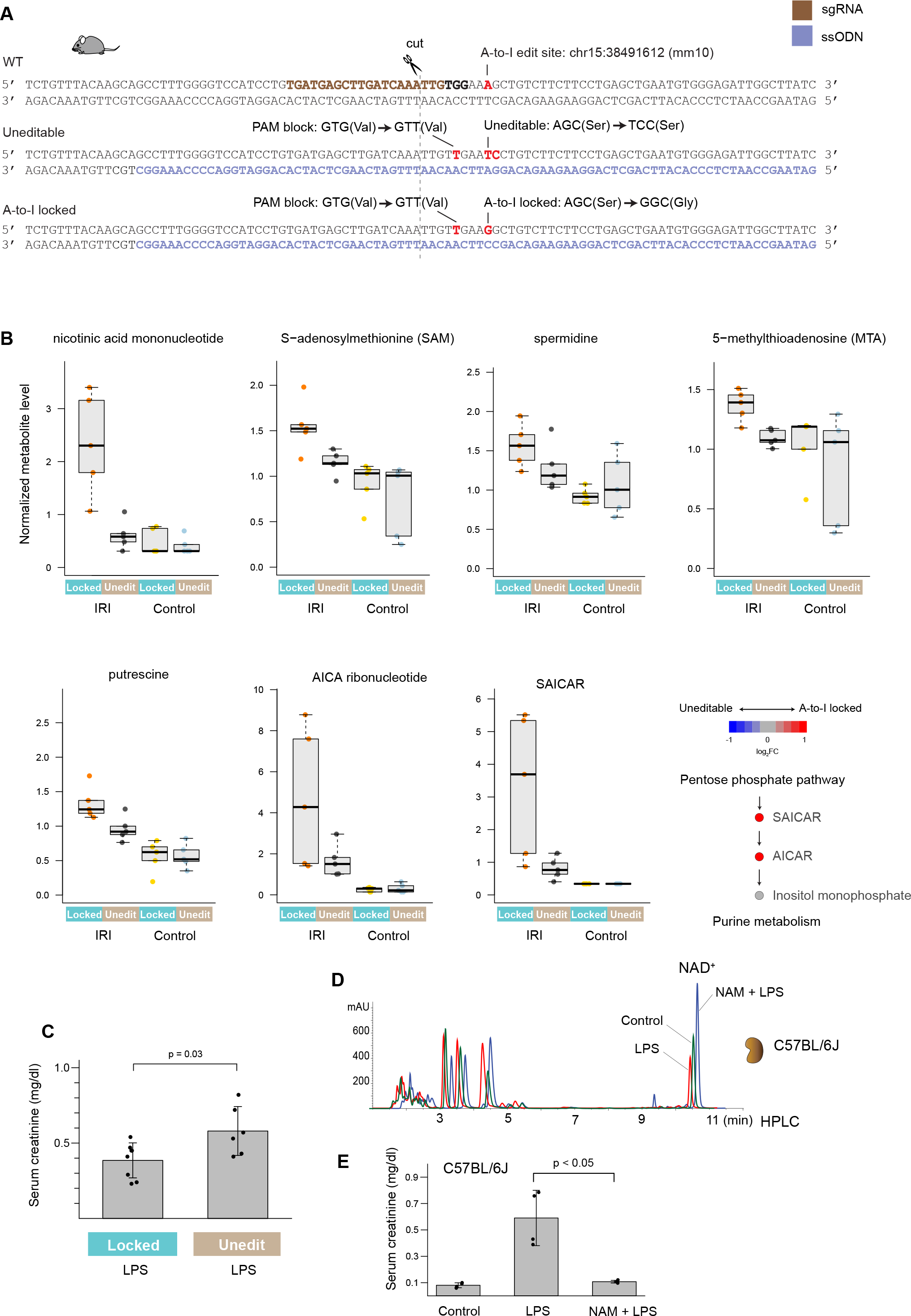
(**A**) CRISPR knock-in design used for generating the Azin1 A-to-I locked and uneditable mice. The single guide RNA is shown in brown, and single-stranded oligonucleotides are shown in blue. Key mutations introduced are annotated. (**B**) Select examples of metabolite levels under the indicated conditions. (**C**) Serum creatinine levels 24 hours after 4 mg/kg LPS intravenously for homozygous Azin1 A-to-I locked and uneditable mice. (**D**) Measurements of NAD^+^ levels in the kidney (C57BL/6J wild-type mice, not Azin1 mutant) by HPLC under the indicated conditions. For clarify, HPLC chromatograms are slightly shifted on the x-axis (elution time). Tissues were harvested 24 hours after 4 mg/kg LPS injection. Supplementation of nicotinamide (NAM, 400 mg/kg intraperitoneally) was performed 1 hour prior to LPS injection. (**E**) Serum creatinine levels under the indicated conditions (C57BL/6J wild-type mice, not Azin1 mutants).

**Supplemental Figure 5.**
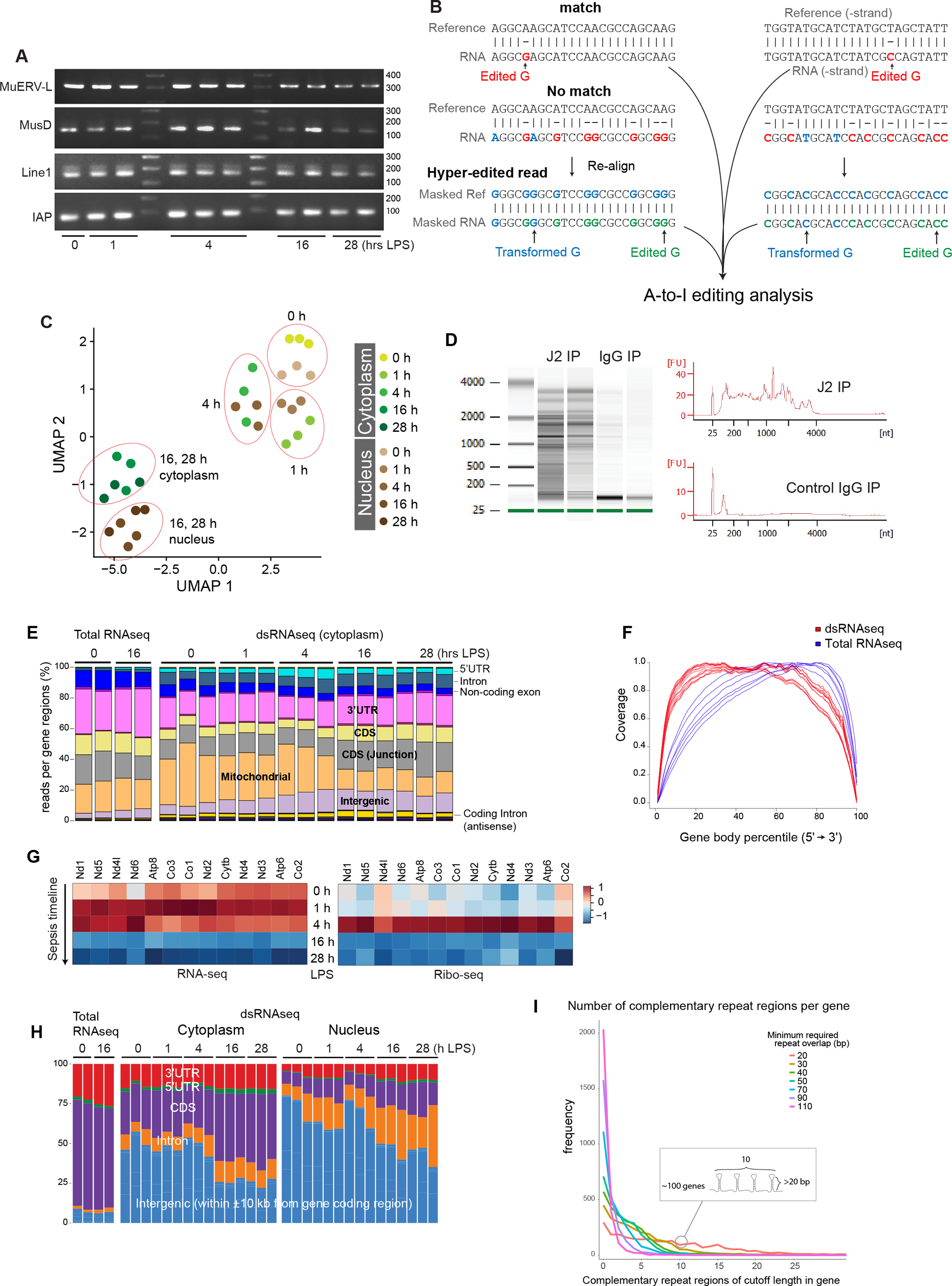
(**A**) PCR gel electrophoresis of select retrotransposons in the kidney under the indicated conditions. Endogenous retroviral elements: Murine endogenous retrovirus-L (MuERV-L), MusD, and Intracisternal A particle (IAP). Long-interspersed nucleotide elements (LINEs): Line1. (**B**) Hyper-editing site alignment strategy, adopted from Porath et al (PMID: 25158696), is shown. Hyper-editing sites were identified by converting all As to Gs in both the genome reference and initially unmapped RNA reads. For clarity, stranded paired-end reads mapped to the minus strand reference is shown as T-to-C mismatches. (**C**) Distribution of dsRNA-seq samples in the UMAP space. (**D**) Representative reads length distribution after J2 dsRNA immunoprecipitation and control IgG2a kappa isotype incubation (Agilent bioanalyzer). (**E**) Distribution of sequence reads mapped to the indicated genome features. (**F**) Distribution of sequence reads mapped to the gene body. The skewed distribution of reads toward the end of gene body in total RNAseq is due to the following: 1. the average length of the last exon is longer than that of the others and 2. there are minimal intronic reads. (**G**) Heatmap displaying transcription and translation levels of mitochondrially-encoded mitochondrial genes over the course of endotoxemia in the kidney. (Reanalysis of published data GEO GSE120877) (**H**) Distribution of sequence reads mapped to the gene coding regions and within ±10 kb from transcription start and end sites. (**I**) Histogram summarizing the frequency of mouse genes harboring complementary repeat regions, dissected based on the lengths of palindromic sequences. This computational search was performed on the reference genome, not on our sequenced files. The x-axis denotes the number of complementary repeat regions within a given gene. The y-axis denotes the number of genes harboring complementary repeat regions.

**Supplemental Figure 6.**
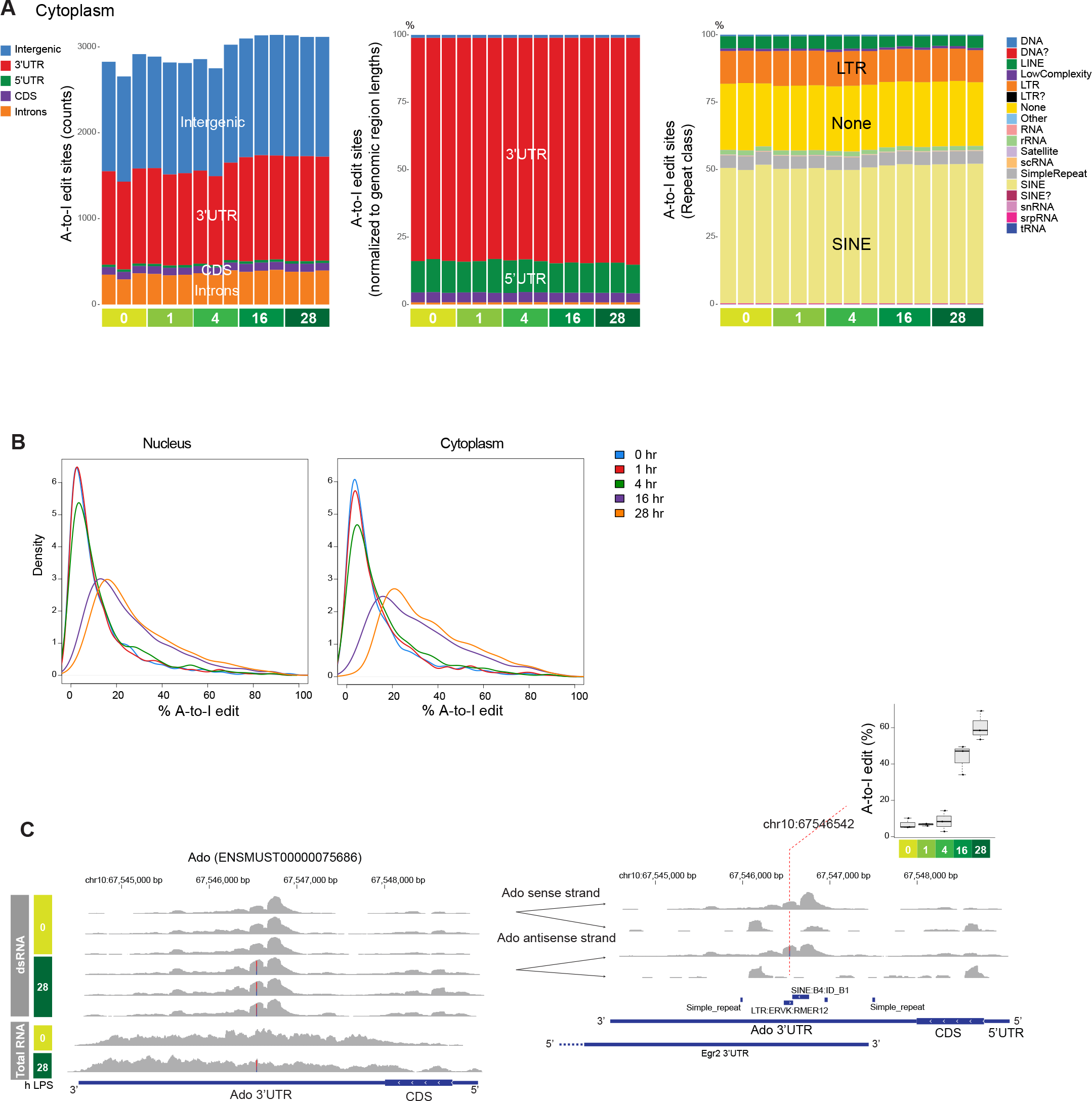
(**A**) Left: Total counts and distribution of A-to-I editing sites per sample (cytoplasmic fraction; editing rate > 10% and reads count > 5 in at least 3 samples; refer to Methods for further preprocessing criteria). Middle: Distribution of A-to-I editing sites, normalized to genomic region lengths. Right: Distribution of A-to-I editing sites per repeat class. (**B**) Density plot displaying A-to-I edit percentages, split by time points and compartments. (**C**) Example of read coverage for the gene Ado with dsRNA enrichment (top 6 tracks) and without dsRNA enrichment (bottom 2 tracks, regular total RNA sequencing). Since the Ado 3’UTR (- strand gene) overlaps with the Egr2 3’UTR (+ strand gene), reads are split into sense and antisense strands to clarify the directionality of A-to-I editing in the right panel. The upper right inset shows A-to-I editing rates at each time point for dsRNA sequencing.

**Supplemental Figure 7.**
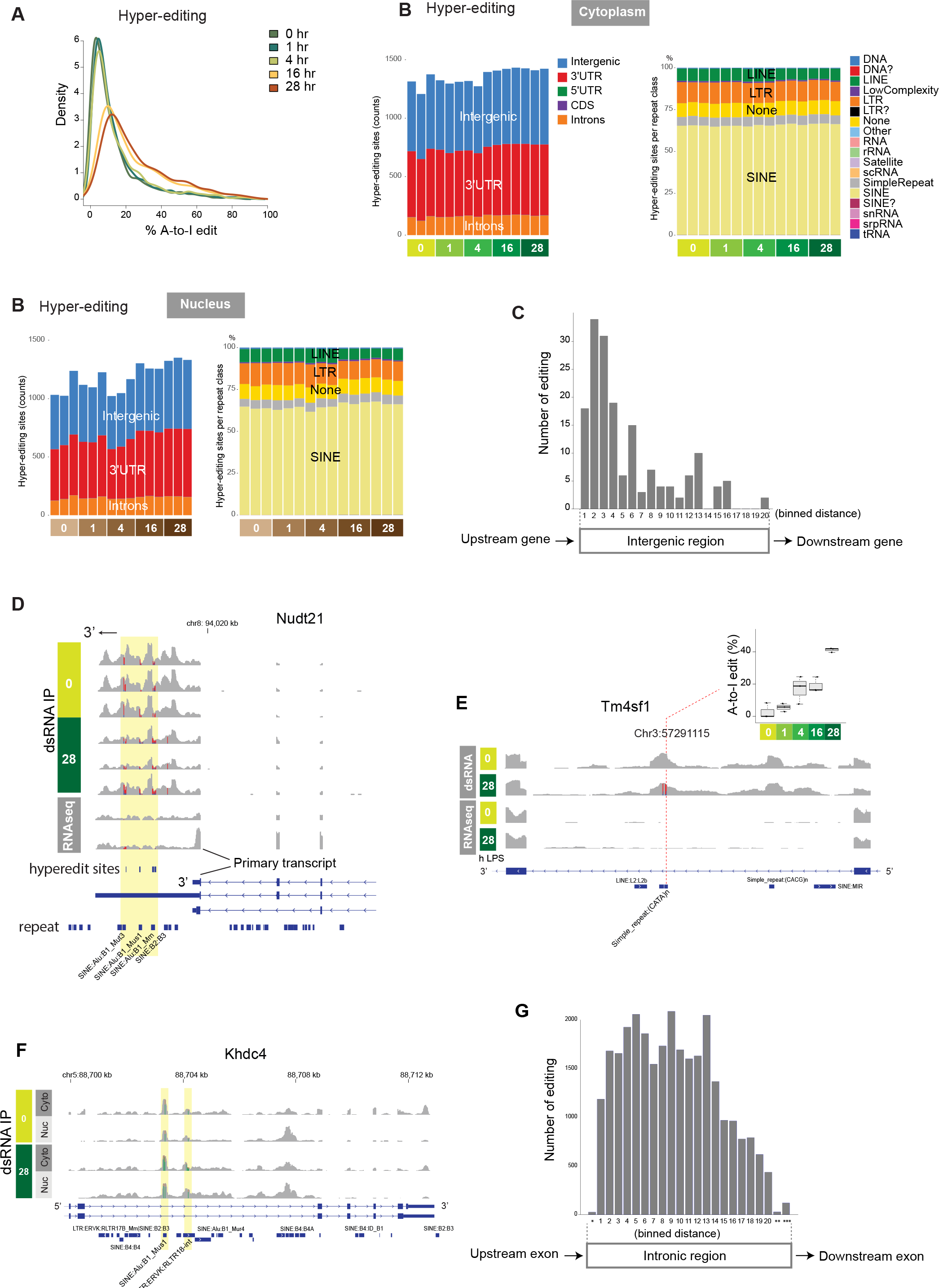
(**A**) Density plot displaying A-to-I edit percentages of hyper-editing sites, split by time points and compartments. (**B**) Summary of hyper-editing sites (cytoplasm in the left two panels and nucleus in the right two panels). (**C**) Distribution of A-to-I editing between gene coding regions. For clarity, regions flanked by genes in the same orientation are shown. (**D**) Example of the enrichment of A-to-I edited reads after the transcription termination site of a primary transcript isoform. The top 6 tracks are dsRNA IP samples, and the bottom 2 tracks are regular total RNA sequencing (0 and 28 hours post LPS). (**E**) Example of read coverage comparison between dsRNA enrichment (top 2 tracks) and without dsRNA enrichment (bottom 2 tracks, regular total RNA sequencing). The inset shows A-to-I editing rates at each time point after endotoxin for dsRNA sequencing. dsRNA immunoprecipitation enriched intronic reads of Tm4sf1, a tetraspanin gene, primarily in the repeat regions, both at 0-hour baseline and later time points. Significant A-to-I editing was observed only at later time points in the endotoxemia model. (**F**) Example of read coverage comparison between nuclear and cytoplasmic fractions. Khdc4, an RNA-binding protein involved in splicing, has several A-to-I editing sites in the intronic region, corresponding to repeat regions. dsRNA immunoprecipitation enriched repeat regions in both nuclear and cytoplasmic fractions, but to a greater extent in the nuclear fraction. (**G**) Distribution of A-to-I editing sites within intronic regions. This intron analysis was done based on sites called by REDItools without further filtration steps, due to limited read coverage depth. *Splice donor site (+6 nucleotides from a splice donor site), **branch point (−40 to −10 nucleotides from a splice acceptor site). ***Splice acceptor site (−10 nucleotides from a splice acceptor site).

**Supplemental Figure 8.**
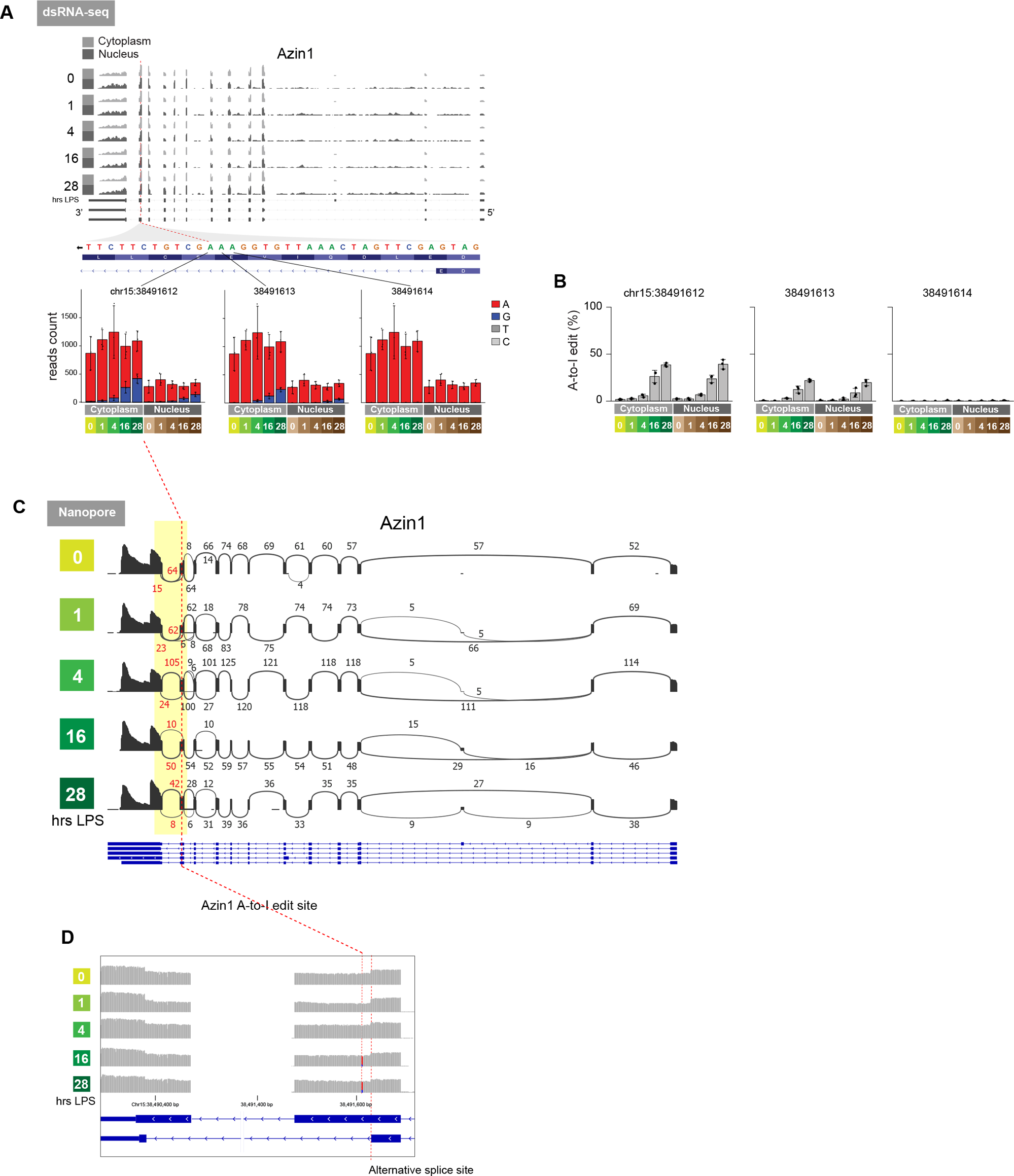
(**A**) Azin1 reads distribution and A-to-I editing under the indicated conditions (dsRNA-seq data). (**B**) Percentage of Azin1 A-to-I editing. (**C**) Sashimi plot displaying reads connectivity for Azin1 under the indicated conditions (Nanopore long-read cDNA sequencing). Approximately 20% of reads near the Azin1 A-to-I editing site exhibit alternative splicing, irrespective of endotoxin time points. (**D**) Magnified view of read coverage near the Azin1 A-to-I editing site and alternative splice site.

## References

1. L. Miller-Fleming, V. Olin-Sandoval, K. Campbell, M. Ralser, Remaining Mysteries of Molecular Biology: The Role of Polyamines in the Cell. J Mol Biol 427, 3389–3406 (2015).

2. F. Madeo, T. Eisenberg, F. Pietrocola, G. Kroemer, Spermidine in health and disease. Science 359, (2018).

3. A. E. Pegg, Functions of Polyamines in Mammals. J Biol Chem 291, 14904–14912 (2016).

4. K. Igarashi, K. Kashiwagi, Modulation of cellular function by polyamines. Int J Biochem Cell Biol 42, 39–51 (2010).

5. H. L. Lightfoot, J. Hall, Endogenous polyamine function--the RNA perspective. Nucleic Acids Res 42, 11275–11290 (2014).

6. S. Mandal, A. Mandal, H. E. Johansson, A. V. Orjalo, M. H. Park, Depletion of cellular polyamines, spermidine and spermine, causes a total arrest in translation and growth in mammalian cells. Proc Natl Acad Sci U S A 110, 2169–2174 (2013).

7. R. A. Casero, Jr., T. Murray Stewart, A. E. Pegg, Polyamine metabolism and cancer: treatments, challenges and opportunities. Nat Rev Cancer 18, 681–695 (2018).

8. M. Al-Habsi, K. Chamoto, K. Matsumoto, N. Nomura, B. Zhang, Y. Sugiura, K. Sonomura, A. Maharani, Y. Nakajima, Y. Wu, Y. Nomura, R. Menzies, M. Tajima, K. Kitaoka, Y. Haku, S. Delghandi, K. Yurimoto, F. Matsuda, S. Iwata, T. Ogura, S. Fagarasan, T. Honjo, Spermidine activates mitochondrial trifunctional protein and improves antitumor immunity in mice. Science 378, eabj3510 (2022).

9. D. J. Puleston, F. Baixauli, D. E. Sanin, J. Edwards-Hicks, M. Villa, A. M. Kabat, M. M. Kaminski, M. Stanckzak, H. J. Weiss, K. M. Grzes, K. Piletic, C. S. Field, M. Corrado, F. Haessler, C. Wang, Y. Musa, L. Schimmelpfennig, L. Flachsmann, G. Mittler, N. Yosef, V. K. Kuchroo, J. M. Buescher, S. Balabanov, E. J. Pearce, D. R. Green, E. L. Pearce, Polyamine metabolism is a central determinant of helper T cell lineage fidelity. Cell 184, 4186–4202 e4120 (2021).

10. W. Slotkin, K. Nishikura, Adenosine-to-inosine RNA editing and human disease. Genome Med 5, 105 (2013).

11. Z. Tian, M. Liang, Renal metabolism and hypertension. Nat Commun 12, 963 (2021).

12. T. Sieckmann, G. Schley, N. Ogel, S. Kelterborn, F. J. Boivin, M. Fahling, M. I. Ashraf, M. Reichel, E. Vigolo, A. Hartner, F. B. Lichtenberger, T. Breiderhoff, F. Knauf, C. Rosenberger, F. Aigner, K. Schmidt-Ott, H. Scholz, K. M. Kirschner, Strikingly conserved gene expression changes of polyamine regulating enzymes among various forms of acute and chronic kidney injury. Kidney Int 104, 90–107 (2023).

13. R. G. Evans, Maybe the various forms of kidney disease are not so mechanistically different? Kidney Int 104, 31–33 (2023).

14. T. Hato, A. Zollman, Z. Plotkin, T. M. El-Achkar, B. F. Maier, S. L. Pay, S. Dube, P. Cabral, M. Yoshimoto, J. McClintick, P. C. Dagher, Endotoxin Preconditioning Reprograms S1 Tubules and Macrophages to Protect the Kidney. J Am Soc Nephrol 29, 104–117 (2018).

15. N. Melis, I. Rubera, M. Cougnon, S. Giraud, B. Mograbi, A. Belaid, D. F. Pisani, S. M. Huber, S. Lacas-Gervais, K. Fragaki, N. Blondeau, P. Vigne, C. Frelin, T. Hauet, C. Duranton, M. Tauc, Targeting eIF5A Hypusination Prevents Anoxic Cell Death through Mitochondrial Silencing and Improves Kidney Transplant Outcome. J Am Soc Nephrol 28, 811–822 (2017).

16. S. Giraud, T. Kerforne, J. Zely, V. Ameteau, P. Couturier, M. Tauc, T. Hauet, The inhibition of eIF5A hypusination by GC7, a preconditioning protocol to prevent brain death-induced renal injuries in a preclinical porcine kidney transplantation model. Am J Transplant 20, 3326–3340 (2020).

17. K. Zahedi, S. Barone, D. L. Kramer, H. Amlal, L. Alhonen, J. Janne, C. W. Porter, M. Soleimani, The role of spermidine/spermine N1-acetyltransferase in endotoxin-induced acute kidney injury. Am J Physiol Cell Physiol 299, C164–174 (2010).

18. K. Zahedi, S. Barone, M. Soleimani, Polyamine Catabolism in Acute Kidney Injury. Int J Mol Sci 20, (2019).

19. S. Zhu, M. Ashok, J. Li, W. Li, H. Yang, P. Wang, K. J. Tracey, A. E. Sama, H. Wang, Spermine protects mice against lethal sepsis partly by attenuating surrogate inflammatory markers. Mol Med 15, 275–282 (2009).

20. W. Liang, K. Yamahara, C. Hernando-Erhard, S. Lagies, N. Wanner, H. Liang, C. Schell, B. Kammerer, T. B. Huber, T. Bork, A reciprocal regulation of spermidine and autophagy in podocytes maintains the filtration barrier. Kidney Int 98, 1434–1448 (2020).

21. X. Li, X. Zhou, X. Liu, X. Li, X. Jiang, B. Shi, S. Wang, Spermidine protects against acute kidney injury by modulating macrophage NLRP3 inflammasome activation and mitochondrial respiration in an eIF5A hypusination-related pathway. Mol Med 28, 103 (2022).

22. C. Kahana, The antizyme family for regulating polyamines. J Biol Chem 293, 18730–18735 (2018).

23. L. Chen, Y. Li, C. H. Lin, T. H. Chan, R. K. Chow, Y. Song, M. Liu, Y. F. Yuan, L. Fu, K. L. Kong, L. Qi, Y. Li, N. Zhang, A. H. Tong, D. L. Kwong, K. Man, C. M. Lo, S. Lok, D. G. Tenen, X. Y. Guan, Recoding RNA editing of AZIN1 predisposes to hepatocellular carcinoma. Nat Med 19, 209–216 (2013).

24. K. Shigeyasu, Y. Okugawa, S. Toden, J. Miyoshi, Y. Toiyama, T. Nagasaka, N. Takahashi, M. Kusunoki, T. Takayama, Y. Yamada, T. Fujiwara, L. Chen, A. Goel, AZIN1 RNA editing confers cancer stemness and enhances oncogenic potential in colorectal cancer. JCI Insight 3, (2018).

25. A. Ghalali, L. Wang, K. H. Stopsack, J. M. Rice, S. Wu, C. L. Wu, B. R. Zetter, M. S. Rogers, AZIN1 RNA editing alters protein interactions, leading to nuclear translocation and worse outcomes in prostate cancer. Exp Mol Med 54, 1713–1726 (2022).

26. Y. R. Qin, J. J. Qiao, T. H. Chan, Y. H. Zhu, F. F. Li, H. Liu, J. Fei, Y. Li, X. Y. Guan, L. Chen, Adenosine-to-inosine RNA editing mediated by ADARs in esophageal squamous cell carcinoma. Cancer Res 74, 840–851 (2014).

27. F. Wang, J. He, S. Liu, A. Gao, L. Yang, G. Sun, W. Ding, C. Y. Li, F. Gou, M. He, F. Wang, X. Wang, X. Zhao, P. Zhu, S. Hao, Y. Ma, H. Cheng, J. Yu, T. Cheng, A comprehensive RNA editome reveals that edited Azin1 partners with DDX1 to enable hematopoietic stem cell differentiation. Blood 138, 1939–1952 (2021).

28. R. Merdler-Rabinowicz, D. Gorelik, J. Park, C. Meydan, J. Foox, M. Karmon, H. S. Roth, R. Cohen-Fultheim, G. Shohat-Ophir, E. Eisenberg, E. Ruppin, C. E. Mason, E. Y. Levanon, Elevated A-to-I RNA editing in COVID-19 infected individuals. NAR Genom Bioinform 5, lqad092 (2023).

29. B. M. Fox, H. W. Gil, L. Kirkbride-Romeo, R. A. Bagchi, S. A. Wennersten, K. R. Haefner, N. I. Skrypnyk, C. N. Brown, D. E. Soranno, K. M. Gist, B. R. Griffin, A. Jovanovich, J. A. Reisz, M. J. Wither, A. D’Alessandro, C. L. Edelstein, N. Clendenen, T. A. McKinsey, C. Altmann, S. Faubel, Metabolomics assessment reveals oxidative stress and altered energy production in the heart after ischemic acute kidney injury in mice. Kidney Int 95, 590–610 (2019).

30. M. Cougnon, R. Carcy, N. Melis, I. Rubera, C. Duranton, K. Dumas, J. F. Tanti, C. Pons, N. Soubeiran, M. Shkreli, T. Hauet, L. Pellerin, S. Giraud, N. Blondeau, M. Tauc, D. F. Pisani, Inhibition of eIF5A hypusination reprogrammes metabolism and glucose handling in mouse kidney. Cell Death Dis 12, 283 (2021).

31. U. B. Gunnia, P. S. Amenta, J. R. Seibold, T. J. Thomas, Successful treatment of lupus nephritis in MRL-lpr/lpr mice by inhibiting ornithine decarboxylase. Kidney Int 39, 882–890 (1991).

32. S. C. Thomson, A. Deng, D. Bao, J. Satriano, R. C. Blantz, V. Vallon, Ornithine decarboxylase, kidney size, and the tubular hypothesis of glomerular hyperfiltration in experimental diabetes. J Clin Invest 107, 217–224 (2001).

33. T. M. Tran, R. Guha, S. Portugal, J. Skinner, A. Ongoiba, J. Bhardwaj, M. Jones, J. Moebius, P. Venepally, S. Doumbo, E. A. DeRiso, S. Li, K. Vijayan, S. L. Anzick, G. T. Hart, E. M. O’Connell, O. K. Doumbo, A. Kaushansky, G. Alter, P. L. Felgner, H. Lorenzi, K. Kayentao, B. Traore, E. F. Kirkness, P. D. Crompton, A Molecular Signature in Blood Reveals a Role for p53 in Regulating Malaria-Induced Inflammation. Immunity 51, 750–765 e710 (2019).

34. E. Picardi, A. M. D’Erchia, C. Lo Giudice, G. Pesole, REDIportal: a comprehensive database of A-to-I RNA editing events in humans. Nucleic Acids Res 45, D750–D757 (2017).

35. B. B. Lake, R. Menon, S. Winfree, Q. Hu, R. M. Ferreira, K. Kalhor, D. Barwinska, E. A. Otto, M. Ferkowicz, D. Diep, N. Plongthongkum, A. Knoten, S. Urata, L. H. Mariani, A. S. Naik, S. Eddy, B. Zhang, Y. Wu, D. Salamon, J. C. Williams, X. Wang, K. S. Balderrama, P. J. Hoover, E. Murray, J. L. Marshall, T. Noel, A. Vijayan, A. Hartman, F. Chen, S. S. Waikar, S. E. Rosas, F. P. Wilson, P. M. Palevsky, K. Kiryluk, J. R. Sedor, R. D. Toto, C. R. Parikh, E. H. Kim, R. Satija, A. Greka, E. Z. Macosko, P. V. Kharchenko, J. P. Gaut, J. B. Hodgin, K. Consortium, M. T. Eadon, P. C. Dagher, T. M. El-Achkar, K. Zhang, M. Kretzler, S. Jain, An atlas of healthy and injured cell states and niches in the human kidney. Nature 619, 585–594 (2023).

36. J. Hansen, R. Sealfon, R. Menon, M. T. Eadon, B. B. Lake, B. Steck, K. Anjani, S. Parikh, T. K. Sigdel, G. Zhang, D. Velickovic, D. Barwinska, T. Alexandrov, D. Dobi, P. Rashmi, E. A. Otto, M. Rivera, M. P. Rose, C. R. Anderton, J. P. Shapiro, A. Pamreddy, S. Winfree, Y. Xiong, Y. He, I. H. de Boer, J. B. Hodgin, L. Barisoni, A. S. Naik, K. Sharma, M. M. Sarwal, K. Zhang, J. Himmelfarb, B. Rovin, T. M. El-Achkar, Z. Laszik, J. C. He, P. C. Dagher, M. T. Valerius, S. Jain, L. M. Satlin, O. G. Troyanskaya, M. Kretzler, R. Iyengar, E. U. Azeloglu, P. Kidney Precision Medicine, A reference tissue atlas for the human kidney. Sci Adv 8, eabn4965 (2022).

37. B. G. Hale, Antiviral immunity triggered by infection-induced host transposable elements. Curr Opin Virol 52, 211–216 (2022).

38. T. Hato, B. Maier, F. Syed, J. Myslinski, A. Zollman, Z. Plotkin, M. T. Eadon, P. C. Dagher, Bacterial sepsis triggers an antiviral response that causes translation shutdown. J Clin Invest 129, 296–309 (2019).

39. D. Janosevic, J. Myslinski, T. W. McCarthy, A. Zollman, F. Syed, X. Xuei, H. Gao, Y. L. Liu, K. S. Collins, Y. H. Cheng, S. Winfree, T. M. El-Achkar, B. Maier, R. Melo Ferreira, M. T. Eadon, T. Hato, P. C. Dagher, The orchestrated cellular and molecular responses of the kidney to endotoxin define a precise sepsis timeline. Elife 10, (2021).

40. A. Kidwell, S. P. S. Yadav, B. Maier, A. Zollman, K. Ni, A. Halim, D. Janosevic, J. Myslinski, F. Syed, L. Zeng, A. B. Waffo, K. Banno, X. Xuei, E. H. Doud, P. C. Dagher, T. Hato, Translation Rescue by Targeting Ppp1r15a through Its Upstream Open Reading Frame in Sepsis-Induced Acute Kidney Injury in a Murine Model. J Am Soc Nephrol 34, 220–240 (2023).

41. L. Li, Y. Mao, L. Zhao, L. Li, J. Wu, M. Zhao, W. Du, L. Yu, P. Jiang, p53 regulation of ammonia metabolism through urea cycle controls polyamine biosynthesis. Nature 567, 253–256 (2019).

42. H. Chen, T. Tong, S. Y. Lu, L. Ji, B. Xuan, G. Zhao, Y. Yan, L. Song, L. Zhao, Y. Xie, X. Leng, X. Zhang, Y. Cui, X. Chen, H. Xiong, T. Yu, X. Li, T. Sun, Z. Wang, J. Chen, Y. X. Chen, J. Hong, J. Y. Fang, Urea cycle activation triggered by host-microbiota maladaptation driving colorectal tumorigenesis. Cell Metab 35, 651–666 e657 (2023).

43. P. Penttinen, J. Jaehrling, A. E. Damdimopoulos, J. Inzunza, J. G. Lemmen, P. van der Saag, K. Pettersson, G. Gauglitz, S. Makela, I. Pongratz, Diet-derived polyphenol metabolite enterolactone is a tissue-specific estrogen receptor activator. Endocrinology 148, 4875–4886 (2007).

44. I. Asplin, G. Galasko, J. Larner, chiro-inositol deficiency and insulin resistance: a comparison of the chiro-inositol- and the myo-inositol-containing insulin mediators isolated from urine, hemodialysate, and muscle of control and type II diabetic subjects. Proc Natl Acad Sci U S A 90, 5924–5928 (1993).

45. P. McLean, S. Kunjara, A. L. Greenbaum, K. Gumaa, J. Lopez-Prados, M. Martin-Lomas, T. W. Rademacher, Reciprocal control of pyruvate dehydrogenase kinase and phosphatase by inositol phosphoglycans. Dynamic state set by “push-pull” system. J Biol Chem 283, 33428–33436 (2008).

46. J. H. Jeon, T. Thoudam, E. J. Choi, M. J. Kim, R. A. Harris, I. K. Lee, Loss of metabolic flexibility as a result of overexpression of pyruvate dehydrogenase kinases in muscle, liver and the immune system: Therapeutic targets in metabolic diseases. J Diabetes Investig 12, 21–31 (2021).

47. K. M. Ralto, E. P. Rhee, S. M. Parikh, NAD(+) homeostasis in renal health and disease. Nat Rev Nephrol 16, 99–111 (2020).

48. G. R. Steinberg, D. G. Hardie, New insights into activation and function of the AMPK. Nat Rev Mol Cell Biol 24, 255–272 (2023).

49. D. P. Reich, B. L. Bass, Mapping the dsRNA World. Cold Spring Harb Perspect Biol 11, (2019).

50. N. Bannert, R. Kurth, Retroelements and the human genome: new perspectives on an old relation. Proc Natl Acad Sci U S A 101 Suppl 2, 14572–14579 (2004).

51. J. Schonborn, J. Oberstrass, E. Breyel, J. Tittgen, J. Schumacher, N. Lukacs, Monoclonal antibodies to double-stranded RNA as probes of RNA structure in crude nucleic acid extracts. Nucleic Acids Res 19, 2993–3000 (1991).

52. A. Dhir, S. Dhir, L. S. Borowski, L. Jimenez, M. Teitell, A. Rotig, Y. J. Crow, G. I. Rice, D. Duffy, C. Tamby, T. Nojima, A. Munnich, M. Schiff, C. R. de Almeida, J. Rehwinkel, A. Dziembowski, R. J. Szczesny, N. J. Proudfoot, Mitochondrial double-stranded RNA triggers antiviral signalling in humans. Nature 560, 238–242 (2018).

53. S. M. Tan-Wong, S. Dhir, N. J. Proudfoot, R-Loops Promote Antisense Transcription across the Mammalian Genome. Mol Cell 76, 600–616 e606 (2019).

54. A. B. Conley, W. J. Miller, I. K. Jordan, Human cis natural antisense transcripts initiated by transposable elements. Trends Genet 24, 53–56 (2008).

55. H. T. Porath, S. Carmi, E. Y. Levanon, A genome-wide map of hyper-edited RNA reveals numerous new sites. Nat Commun 5, 4726 (2014).

56. Y. E. Hsiao, J. H. Bahn, Y. Yang, X. Lin, S. Tran, E. W. Yang, G. Quinones-Valdez, X. Xiao, RNA editing in nascent RNA affects pre-mRNA splicing. Genome Res 28, 812–823 (2018).

57. J. Ramirez-Moya, C. Miliotis, A. R. Baker, R. I. Gregory, F. J. Slack, P. Santisteban, An ADAR1-dependent RNA editing event in the cyclin-dependent kinase CDK13 promotes thyroid cancer hallmarks. Mol Cancer 20, 115 (2021).

58. K. Nishikura, A-to-I editing of coding and non-coding RNAs by ADARs. Nat Rev Mol Cell Biol 17, 83–96 (2016).

59. Y. Shiromoto, M. Sakurai, M. Minakuchi, K. Ariyoshi, K. Nishikura, ADAR1 RNA editing enzyme regulates R-loop formation and genome stability at telomeres in cancer cells. Nat Commun 12, 1654 (2021).

60. H. Chung, J. J. A. Calis, X. Wu, T. Sun, Y. Yu, S. L. Sarbanes, V. L. Dao Thi, A. R. Shilvock, H. H. Hoffmann, B. R. Rosenberg, C. M. Rice, Human ADAR1 Prevents Endogenous RNA from Triggering Translational Shutdown. Cell 172, 811–824 e814 (2018).

61. J. B. Li, E. Y. Levanon, J. K. Yoon, J. Aach, B. Xie, E. Leproust, K. Zhang, Y. Gao, G. M. Church, Genome-wide identification of human RNA editing sites by parallel DNA capturing and sequencing. Science 324, 1210–1213 (2009).

62. J. M. Eggington, T. Greene, B. L. Bass, Predicting sites of ADAR editing in double-stranded RNA. Nat Commun 2, 319 (2011).

63. A. Uzonyi, R. Nir, O. Shliefer, N. Stern-Ginossar, Y. Antebi, Y. Stelzer, E. Y. Levanon, S. Schwartz, Deciphering the principles of the RNA editing code via large-scale systematic probing. Mol Cell 81, 2374–2387 e2373 (2021).

64. T. Sun, Y. Yu, X. Wu, A. Acevedo, J. D. Luo, J. Wang, W. M. Schneider, B. Hurwitz, B. R. Rosenberg, H. Chung, C. M. Rice, Decoupling expression and editing preferences of ADAR1 p150 and p110 isoforms. Proc Natl Acad Sci U S A 118, (2021).

65. C. X. George, Z. Gan, Y. Liu, C. E. Samuel, Adenosine deaminases acting on RNA, RNA editing, and interferon action. J Interferon Cytokine Res 31, 99–117 (2011).

66. R. Melo Ferreira, A. R. Sabo, S. Winfree, K. S. Collins, D. Janosevic, C. J. Gulbronson, Y. H. Cheng, L. Casbon, D. Barwinska, M. J. Ferkowicz, X. Xuei, C. Zhang, K. W. Dunn, K. J. Kelly, T. A. Sutton, T. Hato, P. C. Dagher, T. M. El-Achkar, M. T. Eadon, Integration of spatial and single cell transcriptomics localizes epithelial-immune cross-talk in kidney injury. JCI Insight, (2021).

67. R. Kalakeche, T. Hato, G. Rhodes, K. W. Dunn, T. M. El-Achkar, Z. Plotkin, R. M. Sandoval, P. C. Dagher, Endotoxin uptake by S1 proximal tubular segment causes oxidative stress in the downstream S2 segment. J Am Soc Nephrol 22, 1505–1516 (2011).

68. T. Hato, S. Winfree, R. Kalakeche, S. Dube, R. Kumar, M. Yoshimoto, Z. Plotkin, P. C. Dagher, The macrophage mediates the renoprotective effects of endotoxin preconditioning. J Am Soc Nephrol 26, 1347–1362 (2015).

69. T. Hato, S. Winfree, R. Day, R. M. Sandoval, B. A. Molitoris, M. C. Yoder, R. C. Wiggins, Y. Zheng, K. W. Dunn, P. C. Dagher, Two-Photon Intravital Fluorescence Lifetime Imaging of the Kidney Reveals Cell-Type Specific Metabolic Signatures. J Am Soc Nephrol 28, 2420–2430 (2017).

70. J. Zhou, A. Abedini, M. Balzer, R. Shrestra, P. Dhillon, H. Liu, H. Hu, K. Susztak, Unified Mouse and Human Kidney Single-Cell Expression Atlas Reveal Commonalities and Differences in Disease States. J Am Soc Nephrol, (2023).

71. C. V. Riella, M. McNulty, G. T. Ribas, C. F. Tattersfield, C. Perez-Gill, F. Eichinger, J. Kelly, J. Chun, B. Subramanian, D. Guizelini, N. Nephrotic Syndrome Study, S. L. Alper, M. R. Pollak, M. G. Sampson, D. J. Friedman, ADAR regulates APOL1 via A-to-I RNA editing by inhibition of MDA5 activation in a paradoxical biological circuit. Proc Natl Acad Sci U S A 119, e2210150119 (2022).

72. M. T. Eadon, T. H. Schwantes-An, C. L. Phillips, A. R. Roberts, C. V. Greene, A. Hallab, K. J. Hart, S. N. Lipp, C. Perez-Ledezma, K. O. Omar, K. J. Kelly, S. M. Moe, P. C. Dagher, T. M. El-Achkar, R. N. Moorthi, Kidney Histopathology and Prediction of Kidney Failure: A Retrospective Cohort Study. Am J Kidney Dis 76, 350–360 (2020).

73. D. Barwinska, T. M. El-Achkar, R. Melo Ferreira, F. Syed, Y. H. Cheng, S. Winfree, M. J. Ferkowicz, T. Hato, K. S. Collins, K. W. Dunn, K. J. Kelly, T. A. Sutton, B. H. Rovin, S. V. Parikh, C. L. Phillips, P. C. Dagher, M. T. Eadon, P. Kidney Precision Medicine, Molecular characterization of the human kidney interstitium in health and disease. Sci Adv 7, (2021).

74. Y. Gao, S. Chen, S. Halene, T. Tebaldi, Transcriptome-wide quantification of double-stranded RNAs in live mouse tissues by dsRIP-Seq. STAR Protoc 2, 100366 (2021).

75. C. Lo Giudice, M. A. Tangaro, G. Pesole, E. Picardi, Investigating RNA editing in deep transcriptome datasets with REDItools and REDIportal. Nat Protoc 15, 1098–1131 (2020).

76. Y. Chen, A. Sim, Y. K. Wan, K. Yeo, J. J. X. Lee, M. H. Ling, M. I. Love, J. Goke, Context-aware transcript quantification from long-read RNA-seq data with Bambu. Nat Methods 20, 1187–1195 (2023).

77. D. Kigami, N. Minami, H. Takayama, H. Imai, MuERV-L is one of the earliest transcribed genes in mouse one-cell embryos. Biol Reprod 68, 651–654 (2003).

78. H. M. Rowe, J. Jakobsson, D. Mesnard, J. Rougemont, S. Reynard, T. Aktas, P. V. Maillard, H. Layard-Liesching, S. Verp, J. Marquis, F. Spitz, D. B. Constam, D. Trono, KAP1 controls endogenous retroviruses in embryonic stem cells. Nature 463, 237–240 (2010).

